# Cooperation between oncogenic Ras and p53 stimulates JAK/STAT non-cell autonomously to promote Ras tumor radioresistance

**DOI:** 10.1101/2020.10.05.327098

**Authors:** Yong-Li Dong, Ganghadara P Valdka, Jin-Yu (Jim) Lu, Vakil Ahmad, Thomas J. Klein, Lu-Fang Liu, Peter M. Glazer, Tian Xu, Chiswili Y. Chabu

## Abstract

Oncogenic RAS mutations are associated with tumor resistance to radiation therapy. The underlying mechanisms remain unclear. Emergent cell-cell interactions in the tumor microenvironment (TME) profoundly influence therapy outcomes. The nature of these interactions and their role in Ras tumor radioresistance remain unclear. We used *Drosophila* oncogenic Ras tissues and human Ras cancer cell radiation models to address these questions. We discovered that cellular response to genotoxic stress cooperates with oncogenic Ras to activate JAK/STAT non-cell autonomously in the TME. JAK/STAT accelerates the growth of the less-damaged Ras tumor cells, leading to rapid tumor recurrence. Specifically, p53 is heterogeneously activated in Ras tumor tissues in response to irradiation. This mosaicism allows high p53-expressing Ras clones to stimulate JAK/STAT cytokines, which activate JAK/STAT in the nearby low p53-expressing surviving Ras clones, leading to robust tumor re-establishment. Blocking any part of this cell-cell communication loop re-sensitizes Ras tumor cells to irradiation. This finding suggests that coupling STAT inhibitors to radiotherapy might improve clinical outcomes for Ras cancer patients.

## Introduction

Oncogenic *Ras* mutations activate a complex network of interacting signals to cause aggressive cancers^1,2^. Gold standard treatment options include radiation therapy and conventional chemotherapies that cause irreversible genomic damage and trigger apoptosis^3^. However, oncogenic Ras mutations enable cancer cells to resist these genotoxic agents and evade cell death^4–13^.

Various mechanisms have been proposed to explain the resistance of *Ras* cancers to treatments, including the presence of cancer stem cells in the tumor microenvironment^5,14–16^. Another view is that therapy-resistant cancer cells possess robust DNA repair mechanisms that curtail the apoptotic effect of the treatment. In many cancers, including lung and colorectal cancers where oncogenic Ras mutations are common, there is an association between polymorphisms in DNA-damage response genes and improved clinical response to genotoxic agents^17–23^. However, cellular responses to DNA damage are complex and include activation of cell-cell interactions that we don’t fully understand^24^. How these nonautonomous DNA-damage effects impact the response of Ras cancers to genotoxic therapies is an underexplored area of research. Animal tumor models provide the advantage of interrogating tumor resistance mechanisms at the tissue level, making it possible to identify novel and broadly applicable biology.

In genetic screens for suppressors of oncogenic Ras (*Ras^V12^*)-mediated tissue overgrowth in *Drosophila^25, 26^*, we isolated genotoxic mutations, including null alleles of the Pax2 transactivation domain-interacting protein coding gene (*ptip*^-/-^). Interestingly, *ptip*^-/-^ inhibits the growth of *Ras^V12^* cells but also triggers the overgrowth of the surrounding tissues. PTIP is essential for maintaining genomic stability under normal conditions and following DNA damage^27–29^. Disruption of the PTIP DNA repair complex causes genomic instability and triggers a DNA damage response that culminates in the activation of p53, which orchestrates DNA repair or triggers apoptosis of the damaged cell^29,30^

It is becoming evident that p53 biology is far more complex than initially thought and that it involves nonautonomous functions that are not well understood^31–34^. We found that *ptip*^-/-^ causes genomic instability and, consequently, upregulates p53 in *Ras^V12^* cells. This upregulation of wildtype p53 cooperates with oncogenic Ras signaling to stimulate the secretion of JAK/STAT (Janus kinases/signal transducers and activators of transcription) ligands (interleukin 6-related cytokines known as Unpaired in *Drosophila*). This activates JAK/STAT in the surrounding cells, leading to tissue overgrowth. Ionizing radiation (IR) of *Ras^V12^* tissues or of human Ras cancer cells triggers similar nonautonomous effects. Blocking any part of this p53/*Ras^V12^*-STAT signaling relay inhibits the nonautonomous growth effect and re-sensitizes *Ras^V12^* tissues to IR treatment.

In addition to highlighting the complexity of p53 biology, our work defines a treatment-induced cell-cell interaction dynamic that promotes the recurrence of oncogenic Ras tumors after genotoxic therapies. Our data also provides a possible explanation for why some Ras cancers resist genotoxic therapies despite having no p53 mutations.

## Results

### *ptip*^-/-^ promotes nonautonomous tissue overgrowth

In *Drosophila*, the MARCM (Mosaic Analysis by a Repressible Marker) technique permits the expression of oncogenic Ras (*Ras^V12^*) in clones of cells within the developing eye epithelium^26^. Theses clones co-express a fluorescent protein, making them distinguishable from the surrounding wild-type cells. *Ras^V12^*-mediated tissue overgrowth is readily detectable by the appearance of large and hyperplastic fluorescent clones (tumors) that ultimately kill the animal^35–37^. *Ras^V12^* suppressor mutations are isolated through the identification of mutations that significantly reduce clone size and rescue animal viability when introduced in *Ras^V12^*-expressing cells^35^.

We isolated several *Ras^V12^* suppressors, including mutation *#3804*, using this approach. Mutation *3804* potently suppresses *Ras^V12^*-mediated tumor overgrowth and yields viable adult animals (Fig. 1a versus 1b; 1c versus 1d; 1g-l). To determine whether this mutation synthetically suppresses oncogenic Ras or is cell deleterious on its own, we generated wild-type or *3804* mutant clones in developing and adult eye tissues. Adult eye clones are marked by the absence of the red pigment. As expected, wild-type cells contributed ~50% to the eye field. In contrast, *3804* mutant cells were barely detectable in tissues, arguing that the affected gene is essential for cell viability (Fig. 1m-o).

**Figure 1.**
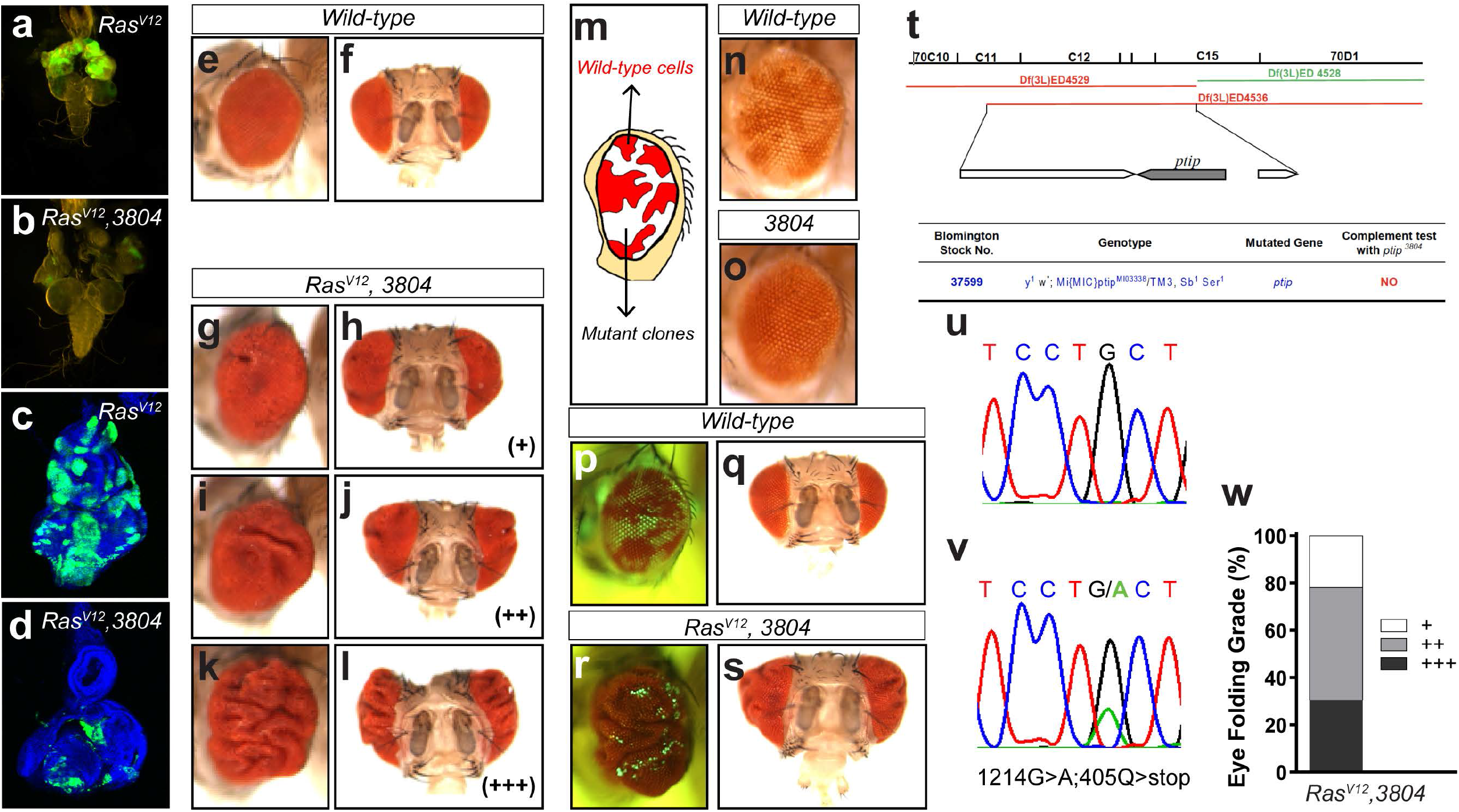
*Ptip*^-/-^ cooperates with oncogenic Ras to induce nonautonomous tissue overgrowth. (a-d) Mutation *3804* suppresses *Ras^V12^*-mediated tumor overgrowth. (a, b) images of third-instar larvae cephalic complexes showing GFP-marked *Ras^V12^* (a) or *Ras^V12^, 3804* (b) clones. Images of eye imaginal discs dissected from third-instar larvae cephalic complexes and containing *Ras^V12^* or *Ras^V12^, 3804* GFP-positive clones are shown in c and d, respectively. (e-l) Side and frontal images of adult fly eyes. The nonautonomous overgrowth phenotype was categorized into three grades based on the severity of the phenotype (+: weak; ++: moderate; +++: strong). (e and f) represent adult eye from wild-type animals. The adult eye tissues bearing *Ras^V12^, 3804* double mutant clones showed varying grades of tissue folding (g-l). (m-o) Mosaic adult eyes bearing wild-type cells and mutant clones marked by a lack of pigmentation. A schematic of the mosaic adult eye is presented in (m). Wild-type cells (n, white color) contribute ~50% to the eye field, whereas the 3804 mutant clones (o) are barely detectable. (p-s) Matched light and fluorescence images of adult eyes containing GFP-positive wild-type (p, q) or *Ras^V12^, 3804* double mutant clones (r, s). (t) Genetic complementation test of the *plip*^-/-^ mutation using overlapping chromosomal deficiency lines. The deficiency line shown in green complements the *ptip*^-/-^ mutation, while the deficiency lines marked in red fail to complement. (u, v) Sequence results showing a G>A mutation in *PTIP*, causing a premature stop sequence. (w) Quantification of (g-l).

Interestingly, although *3804* suppressed *Ras^V12^* tumour overgrowth in the developing eye tissue and correspondingly yielded adult animals, the eyes of the rescued animals were overgrown to varying extents, as evidenced by the appearance of tissue folds (Fig. 1e, f versus 1g-l; w). The overgrown tissues exclusively consisted of wild-type cells (GFP-negative) (Fig. 1p-s).

We used deficiency mapping and allele sequencing approaches and determined that *3804* represents a null mutation in the PAX transcription activation domain interacting protein (PTIP). *3804* animals are homozygous lethal. Two independent deficiency alleles (ED4529 and ED4536) that overlap at the PTIP gene fail to complement *3804* animal lethality (Fig. 1t). Direct sequencing of the *3804* allele revealed a G>A mutation leading to a premature stop codon in the protein (Fig. 1u, v). Thus, we hereafter refer to *3804* as *ptip*^-/-^ and conclude that while *ptip*^-/-^ suppresses growth cell intrinsically, it cooperates with oncogenic Ras to promote the growth of the surrounding tissue.

### Ptip^-/-^ promotes nonautonomous tissue overgrowth via p53

We sought to delineate the underlying mechanism. PTIP was originally identified in a yeast two-hybrid screen^38^. In mammals, PTIP interacts with histone methyltransferase complexes to control developmental transcription programs^38,39^. Also, PTIP is essential for maintaining genomic stability^29,40,41^. Knocking down PTIP causes genomic instability, leading to the activation of p53, which triggers cell cycle arrest or apoptosis^42^. We investigated whether *ptip*^-/-^ causes DNA damage in *Drosophila* epithelial *Ras^V12^* cells and found that it does. DNA damage triggers phosphorylation of a Histone 2A variant (γH2Av), which can be readily detected in immunostaining experiments using phospho-specific antibodies^43,44^. We examined γH2Av in *Ras^V12^* cells in the absence or presence of the *ptip*^-/-^ mutation and detected an increase of γH2Av nuclear foci in the *Ras^V12^ptip*^-/-^ cells compared to the *Ras^v12^* control cells (Fig. 2a-b’). Consistent with DNA damage, *Ras^V12^ptip*^-/-^ double mutant cells had high levels of P53 compared to *Ras^V12^* cells (Fig. 2c-d).

**Figure 2.**
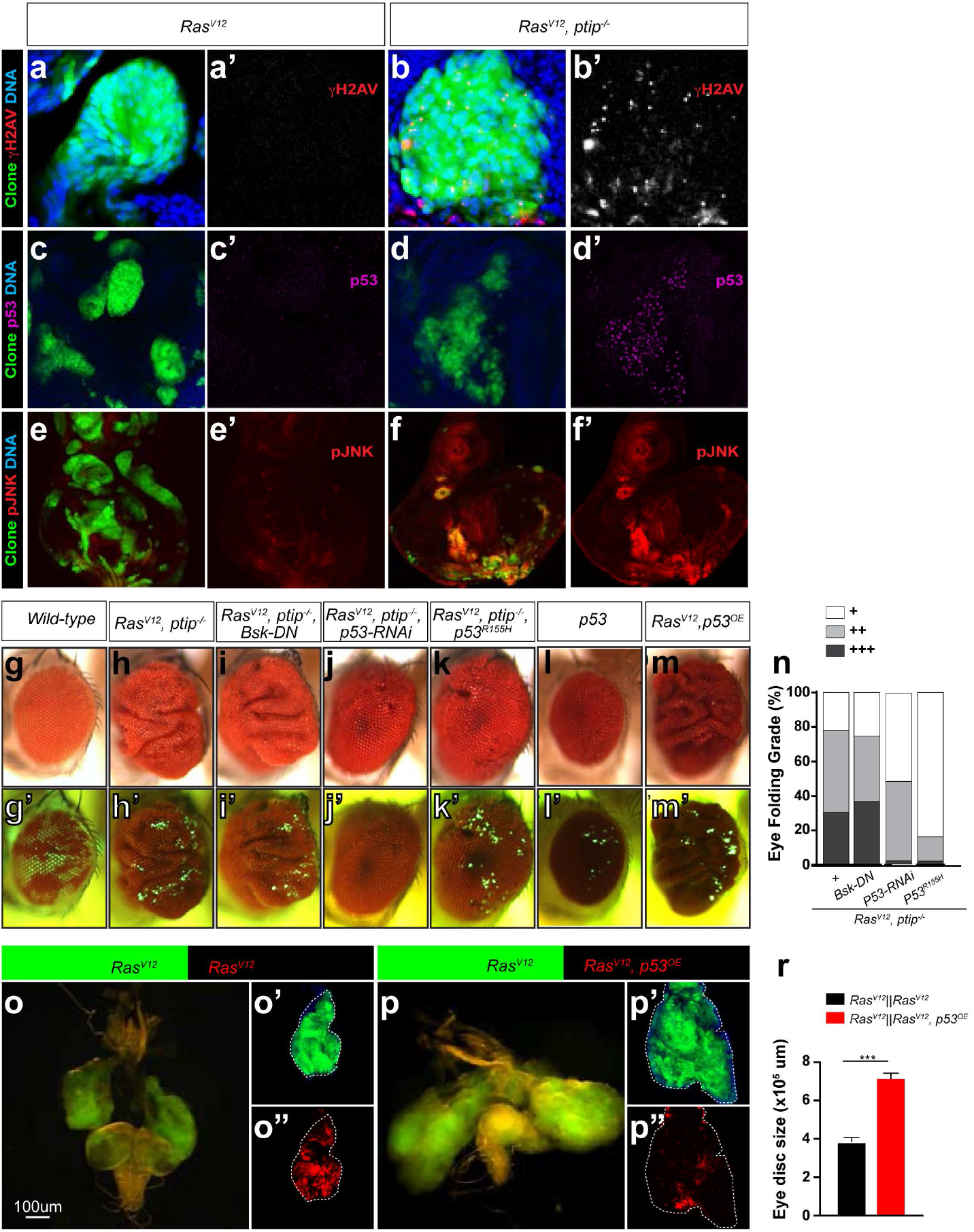
*Ptip*^-/-^ and oncogenic Ras cooperatively induce non-autonomous tissue overgrowth via wild-type p53. (a-f) Representative images of dissected eye imaginal discs containing *Ras^V12^* or *Ras^V12^ptip*^-/-^ double mutant clones (GFP) stained with DAPI to detect DNA or anti-phosphorylated H2AV antibodies to detect DNA damage (a-b’) or anti-p53 (c-d’) or anti-phosphorylated JNK (e-f’) antibodies to detect cellular response to DNA damage. (g-m) Matched light and fluorescence images of adult eyes containing GFP-labelled clones. The respective clone genotypes are indicated at the top of each panel. The corresponding fluorescent images are shown below in (g’-m’). GPF-negative tissues represent wild-type tissues. (n) Quantification of the nonautonomous growth phenotype of adult eyes containing clones of the indicated genotypes: *Ras^V12^ptip^-/-^, Ras^V12^ptip^-/-^ Bsk^DN^, Ras^V12^ptip^-/-^p53^R155H^*, or *Ras^V12^ptip^-/-^ p53^RNAi^*. (o-p) Genetic juxtaposition of GFP-labeled *Ras^V12^* clones with RFP-labeled *Ras^V12^* clones (o-o”, controls) or with RFP-labeled clones of cells co-expressing *Ras^V12^* and wild-type *p53* (*Ras^V12^*, p53^OE^) (p-p”). GFP-positive Ras^V12^ clones are surrounded by RFP/GFP double-positive Ras^V12^ clones (o-o”) or by RFP/GFP double-positive *Ras^V12^, p53^OE^* clones (p-p”). Brain cephalic complex images showing the growth of *Ras^V12^* clones when juxtaposed to *Ras^V12^* or to *Ras^V12^, p53^OE^* clones are shown in o and p, respectively. (r) Quantification of eye tissue sizes from (o-p”). *N= 10* tissues per genotype. Error bars denote SEM values. T-test significance levels is (***) *p*<0.001.

We posited that cellular response to genomic stress likely underlies the nonautonomous growth effect of *ptip*^-/-^ on the *Ras^V12^* clones. The Janus N-terminal Kinase (JNK, also known as Bsk in *Drosophila*) and p53 play central roles in cellular response to DNA damage^45–47^. In addition, we and others have shown that JNK promotes nonautonomous tissue growth in *Drosophila*^35,48,49^, raising the possibility that *ptip*^-/-^ acts via JNK to drive nonautonomous growth in *Ras^V12^* mosaic tissues. Consistent with this, JNK is activated in *Ras^V12^ ptip*^-/-^ mutant cells compared to *Ras^V12^* control cells (Fig. 2e-f’). To directly test this, we inhibited JNK by expressing a potent dominant-negative JNK transgene (*Bsk^DN^*) in *Ras^V12^ptip*^-/-^ cells (*Ras^V12^ptip*^-/-^*Bsk^DN^* triple defective cells) and asked whether this suppresses the nonautonomous tissue overgrowth phenotype. *Bsk^DN^* failed to suppress *Ras^V12^ptip*^-/-^ nonautonomous tissue overgrowth (Fig. 2g-i and 2g’-i’), making it unlikely that JNK plays a significant role in this phenomenon.

We explored alternative mechanisms. Nonautonomous growth-inducing clones (*Ras^V12^ptip*^-/-^ mutant cells) show high levels of wild-type p53 protein (p53^wt^) compared to *Ras^V12^* cells, which do not cause nonautonomous growth (Fig. 2c-d’). Given that the *ptip*^-/-^ mutation occurs very early and that it is permanent, the resulting high p53^wt^ protein levels likely persist throughout the life of the *Ras^V12^* cells. Normally, p53^wt^ has a high turnover rate^50^. We wondered whether the elevated p53^wt^ protein levels observed in the *Ras^V12^ptip*^-/-^ mutant cells play an active role in the nonautonomous tissue growth effect. Indeed, RNAi knockdown of p53 in *Ras^V12^ptip*^-/-^ cells remarkably suppressed the nonautonomous tissue overgrowth (Fig. 2 h, h’, j, j’, and n). Similarly, blocking p53 transcriptional activity in *Ras^V12^ptip*^-/-^ cells by expressing a DNA binding defective p53 mutant version (*p53^R155H^*) also suppressed the overgrowth of the surrounding wild-type cells (Fig. 2 h, h’, k, k’, and n). In addition, direct overexpression of *p53^wt^* (*p53^OE^*) in clones of *Ras^V12^* cells was sufficient to trigger the overgrowth of the surrounding wild-type tissue, mimicking *Ras^V12^ptip*^-/-^ clones (Fig. 2 g, g’, m, m’). The ability of p53^wt^ to drive nonautonomous tissue overgrowth requires oncogenic Ras. Overexpression of p53^wt^ alone failed to generate a similar effect (Fig. 2 g, g’, l, l’).

To assess whether this *Ras^V12^*/p53 cooperation similarly accelerates the growth of adjacent *Ras^V12^* cells, we used the MARCM technique to juxtapose *Ras^V12^p53^OE^* clones (RFP-labelled) with *Ras^V12^* clones (GFP-labelled) (see methods). We found that this alteration caused a massive overgrowth of *Ras^V12^* clones as compared to controls (abutting *Ras^V12^* clones without *P53^OE^*) (Fig. 2o-p” and r). *Ras^V12^ptip*^-/-^ clones had a similar nonautonomous effect on Ras clones (Supplementary Fig. 1). Taken together, these findings indicate that oncogenic Ras cooperates with elevated wild-type p53 protein to drive tumor overgrowth via a novel non-autonomous mechanism.

### Oncogenic Ras and p53 cooperatively stimulate JAK/STAT cytokines to promote nonautonomous tissue overgrowth

We set out to delineate the underlying mechanism. *Drosophila* JAK/STAT ligands Unpaired (Upd, Upd2, and Upd3) mediate non-autonomous tissue growth^35,48,49^. We asked to what extent oncogenic Ras and p53^OE^ act via JAK/STAT. Immunostaining experiments using a *upd* reporter line, *upd-lacZ*^51^, to monitor *upd* transcriptional activity revealed that *p53^OE^* causes *Ras^V12^* cells to upregulate *upd* (Fig. 3a-b’). In a complementary quantitative Polymerase Chain Reaction (qPCR) approach, we found that *p53^OE^* causes *Ras^V12^* cells to upregulate all of the *upd* ligands (upd1-3) in tissues (Fig. 3d). The *ptip*^-/-^ mutation showed similar effects in *Ras^V12^* tissues, including activation of JAK/STAT in cells surrounding the mutant (*Ras^V12^ptip*^-/-^) clones (Fig. 3a, a’, c, c’; Supplementary Fig. 2a-d and 2g-j). These findings suggest that *Ras^V12^p53^OE^* clones induce the growth of surrounding cells via the secretion of JAK/STAT cytokines. To functionally test this, we blocked the secretion of JAK/STAT cytokines to the tissue surrounding *Ras^V12^p53^OE^* clones and asked whether this suppresses the nonautonomous growth effect. We simultaneously knocked down *upd* and *upd2* inside *Ras^V12^p53^OE^* clones by combining *upd-RNAi* expression with *upd2* deletion mutants. This dramatically reduced tissue size (Fig. 3f-i). Knocking down upd in *Ras^V12^ptip*^-/-^ clones similarly suppressed nonautonomous tissue overgrowth (Supplementary Fig. 2e, f, k, and l).

**Figure 3.**
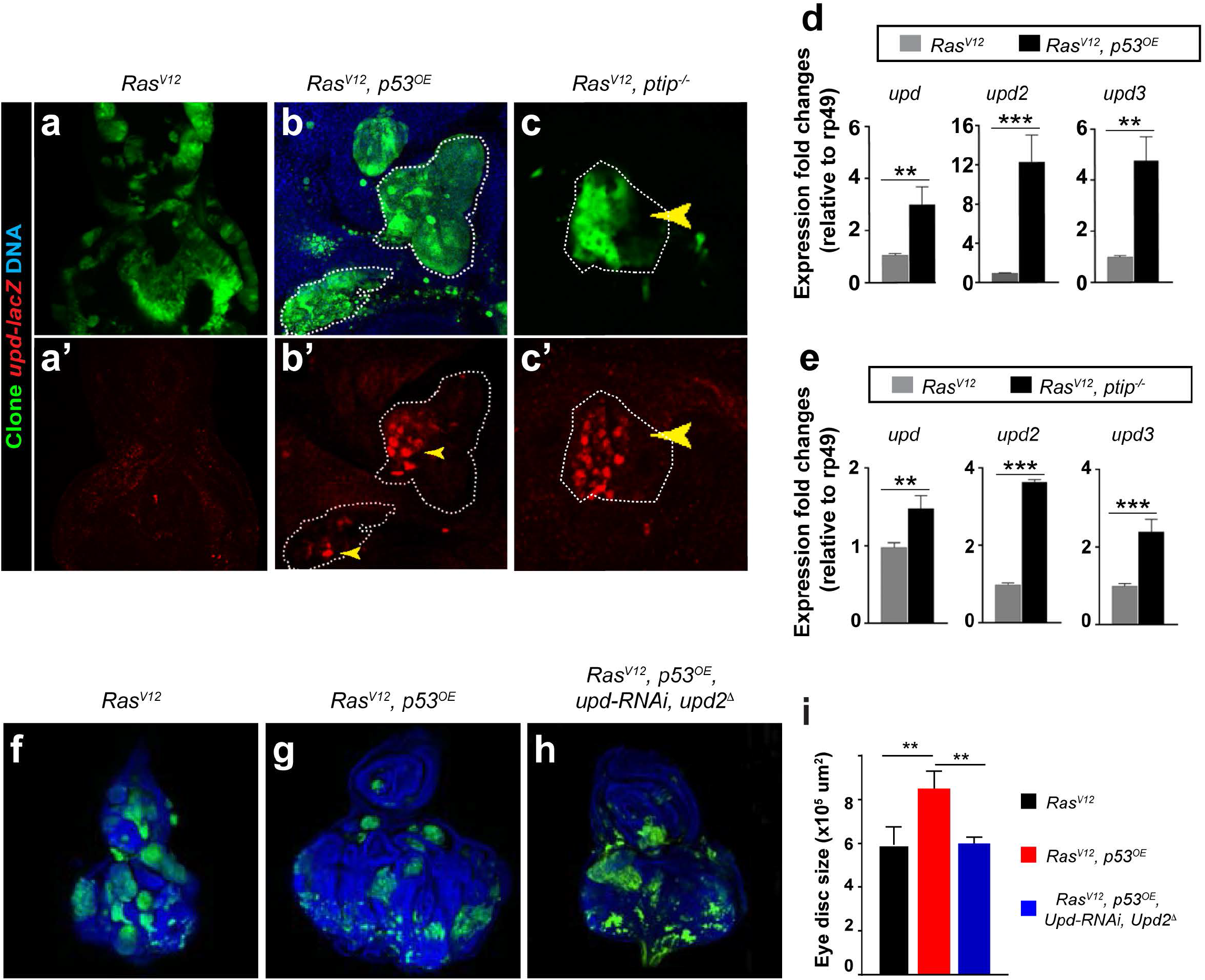
Wild-type *p53* and oncogenic Ras cooperatively stimulate STAT cytokines to drive nonautonomous tissue overgrowth. (a-c’) Images showing *upd-lacZ* expression in eye imaginal discs bearing *Ras^V12^* (a, a’), *Ras^V12^p53^OE^* (b, b’), and *Ras^V12^ptip^-/-^* (c, c’) clones. Anti-*β*gal antibodies were used in immunostaining experiments to detect LacZ. The individual LacZ channel images are shown in a’-c’. The dotted white lines denote clone boundaries and depict representative examples of *upd* overexpression (yellow arrowheads). (d-e) Graphical representation of qPCR data showing expression of *upd, upd2*, and *upd3* in *Ras^V12^* versus *Ras^V12^p53^OE^* (d) or *Ras^V12^* versus *Ras^V12^ptip*^-/-^ (e) eye imaginal disc tissues. Expression was normalized to the transcript abundance of the housekeeping gene *rp49*. T-test significance levels are (**) *p*<0.01 and (***) *p*<0.001. (f-h) Representative images showing overall tissue size of dissected eye imaginal discs harboring GFP-labeled *Ras^V12^* (f) or *Ras^V12^p53^OE^* (g) or *Ras^V12^p53^OE^upd^RNAi^upd2*^Δ^ (h) clones. Tissues were stained with DAPI. (i) Quantification of tissue sizes from (f-h). *N*=10 tissues. Error bars denote SEM. The t-test significance level (**) is *P*<0.01.

We evaluated our findings in breast and lung human cancer cells using supernatant transfer experiments. Breast epithelial cells MCF-10A were cultured in media conditioned with MCF-10A cells (controls) or MCF-10A cells overexpressing wild-type p53 alone or co-expressing oncogenic *HRAS*. STAT signaling was determined in western blot experiments with antibodies that specifically detect activated STAT (anti-phosphorylated STAT). Growth was determined by scoring cell numbers. In this and subsequent experiments, the superscripts “*P53OE*” or “*HRasG12V*” denote overexpression of wild-type *p53* or oncogenic *Ras*, respectively. MCF10A^*HRasG12V, P53OE*^ conditioned media stimulated STAT signaling and correspondingly induced the growth of MCF-10A cells (Fig. 4a, c). This growth promoting effect was significantly reduced when the conditioning cells lacked oncogenic Ras (MCF-10A^*P53OE*^) (Fig. 4c).

**Figure 4.**
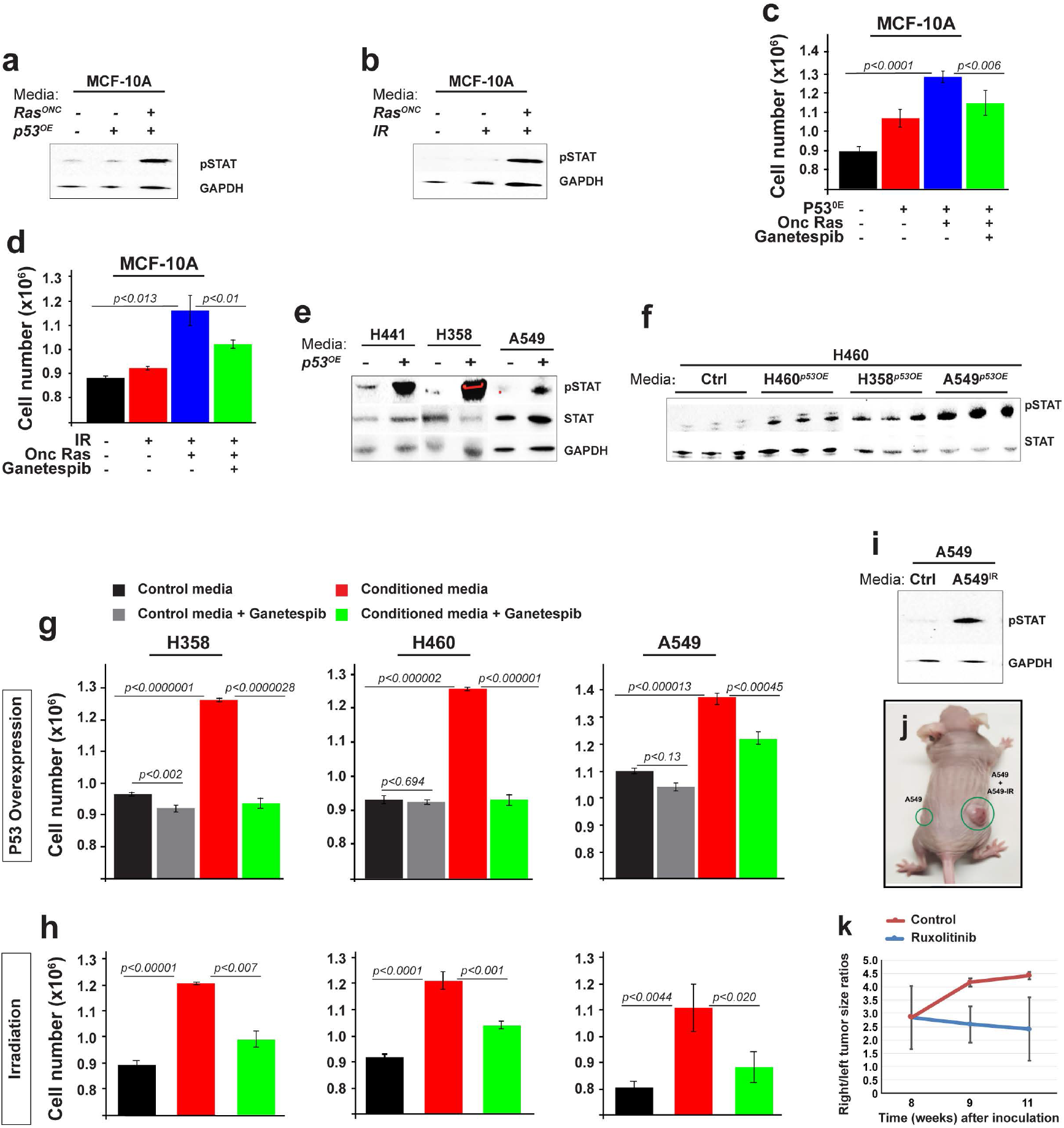
Wild-type p53 and oncogenic Ras paracrine STAT activation stimulates the growth of human cancer cells. (a-i) *In vitro* experiments showing that media conditioned with *p53*-overexpressing or irradiated cancer cells stimulate cell proliferation via STAT signaling. (a, b) Anti-activated STAT (phospho-STAT) and anti-GAPDH (loading control) antibodies Western blots showing STAT activity in MCF-10A cells cultured in media conditioned with MCF-10A (controls) or with MCF-10A cells expressing oncogenic *HRas* (*Ras^ONC^*) or wild-type *p53* (*p53^OE^*) alone or together. (b) Western blot image showing STAT activity in cells conditioned with media conditioned with irradiated MCF-10A cells or MCF-10A cells expressing oncogenic Ras. (c, d) The conditioned media stimulate cell growth, as determined by cell number under the indicated conditions: MCF-10A cells cultured in media conditioned with MCF-10A control cells or MCF-10A cells overexpressing p53 or oncogenic Ras or co-expressing both (c). The growth of MCF-10A cells in media conditioned with irradiated MCF-10A cells in the presence or absence of *p53* overexpression is shown in (d). Ganetespib (25 nm) treatment suppressed cell growth. (e, f). Western blots showing STAT activity in lung cancer cells cultured in media conditioned with unmodified cells or with cells overexpressing *p53*. Total STAT and GAPDH protein levels were used as loading controls. These experiments are shown in biological triplicates in (f) (g) Effect of media conditioned with *p53* overexpressing lung cancer cells on cell number. Ganetespib (25 nm) treatment had no to minimal effect on the growth of control cells but it significantly suppressed the growth of cells growing in conditioned media. (h) Effect of media conditioned with irradiated lung cancer cells on cell number in the presence or absence of Ganetespib (25 nm). (i) Western blot on lysates prepared from A549 lung cancer cells grown in media conditioned with other A549 cells or with irradiated A549 cells. Blots were probed with phosphor-STAT to detect active STAT and GAPDH as a loading control. (j) Image of a nude mouse showing the size of tumor xenographs (green circles) 8 weeks after flank injection of 1×10^6^ A549 cells. Left flank inoculants consisted of normal A549 cells, while right flank received an equal mixture of normal and irradiated A549 cells. (k) Quantification of (j). Tumor sizes were calculated with a digital caliper. To the exception of one animal that developed a larger tumor on the left flank (homogenous inoculants), the remaining five animals developed noticeably larger tumors from the heterogenous inoculants. The graph shows the right to left flank tumor size ratio. Two of the five animals received Ruxolitinib (10 mg/kg) by oral gavage for 3 weeks. The remaining animals were treated with DMSO vehicle control for the same duration. Caliper measurements determined tumor size in treated versus vehicle control animals. Right to left tumor size ratios are shown at 1 and 2 weeks following treatment initiation.

Similar results were observed in lung cancer cells. We overexpressed wild-type p53 in H460, A549, H358, and H441 lung cancer cells, generating H460^*P53OE*^, A549^*P53OE*^, H358^*P53OE*^, and *H441^P53OE^*. Genetically, all of these cells carry oncogenic *Ras* mutations. However, endogenous p53 is wild-type in H460 and A549 and it is mutated in H358 and H441 cells^52^. Compared to matched controls, media conditioned with H460^*P53OE*^, A549^*P53OE*^, H358^*P53OE*^, and H441^*P53OE*^ cells stimulated STAT signaling and cell growth (Fig. 4e-g). Media conditioned with irradiated breast (MCF-10A^*HRasG12V*^) or lung (H358, H460, and A549) cancer cells generated similar nonautonomous effects (Fig. 4b, d, h, and i). Importantly, blocking STAT signaling with the small molecule inhibitor Ganetespib^53^ suppressed the nonautonomous growth-inducing effect of p53^OE^ and IR in both breast and lung cancer cells (Fig. 4c, d, g, and h).

Moreover, we tested our findings *in vivo* using mouse xenograph experiments. We inoculated a total of one million A549 cells into each of the flanks of nude mice. Left flank inoculants were unmodified, while right flanks inoculants consisted of a 50-50 mixture of untreated and irradiated cells. Eight weeks following treatment, tumor xenographs from the mixed population (right flanks) grew markedly larger than those arising from the homogenous inoculants (left flanks) in the same animal (Fig. 4j, k; 83%; *N=6* animals). To test whether STAT plays a role in the observed tumor overgrowth, as suggested by our tissue culture and fly data, we treated animals with the validated pharmacological STAT blocker Ruxolitinib^54^. Animals were treated orally with Ruxolitinib at 10 mg/kg, a dose that is well-tolerated in nude mice^57^. We found that Ruxolitinib suppressed the overgrowth of tumors arising from the mixed cells (Fig. 4k).

JAK/STAT signalling supports tissue growth by promoting cell survival or cell proliferation^55,56^. To distinguish between these two mechanisms, we examined cell death and cell proliferation in the human cells using flow cytometry approaches. We found that p53^OE^ and IR-induced nonautonomous JAK/STAT mainly stimulates cell proliferation (Supplementary Fig. 3 and Supplementary Fig. 4)

Taken together, the above data indicate that stimulation of wild-type p53 cooperates with oncogenic Ras to induce JAK/STAT in the surrounding cells, resulting in nonautonomous growth.

### *The Oncogenic Ras/p53-STAT* signal relay promotes the radioresistance of Drosophila *Ras^V12^* tumor tissues

In several cancer types, oncogenic Ras mutations are associated with disease recurrence following radiation therapy and genotoxic chemotherapies^4–13^.

We used *Drosophila Ras^V12^* tumor tissues under ionizing radiation (IR) conditions to test whether IR-stimulated p53 generates similar effects in a tissue context. Specifically, we asked whether p53 cooperates with oncogenic Ras to establish tumor recurrence via the nonautonomous STAT signal relay described above. We define tumor radioresistance as incomplete sensitivity and/or tumor capacity to rapidly reform following IR treatments.

We first performed a dose finding study to determine a dose that generates cellular effects without grossly impeding animal development. Second-instar larvae harboring GFP-labeled oncogenic Ras clones in eye imaginal discs were treated with 600R or 1000R or 2000R. Each dose was administered three times in 6 hour intervals to mimic clinical settings where total radiation treatments are administered in fractions^57,58^. We found that larvae treated with the 3 x 600R dose developed normally into adults without any detectable abnormalities and those treated with 3 x 2000 died during the pupal stage. Larvae that received 3 x 1000R yielded adult flies with mild rough eyes (data not shown), making it an ideal dosing regimen for our study.

We next asked to what extent IR recapitulates fundamental aspects of the *Ras^V12^/p53^OE^-STAT* signal relay: that is, stimulation of both p53 and JAK/STAT cytokines. Compared to non-treated *Ras^V12^* control tissues, IR elevated p53 protein levels in the immunostaining experiments. Note that p53 stimulation is non-uniform, and p53 was undetectable in portions of wild-type (GFP-negative) and *Ras^V12^* cells (GFP-positive) (Fig. 5a-b’). This mosaicism supports our finding that *Ras^V12^* cells with high p53 protein levels stimulate the growth of the surrounding *Ras^V12^* cells with lower p53 levels (Fig. 2k-l’). qPCR experiments showed that IR transcriptionally stimulates all unpaired cytokines (upd1-3) (Fig. 5c).

**Figure 5.**
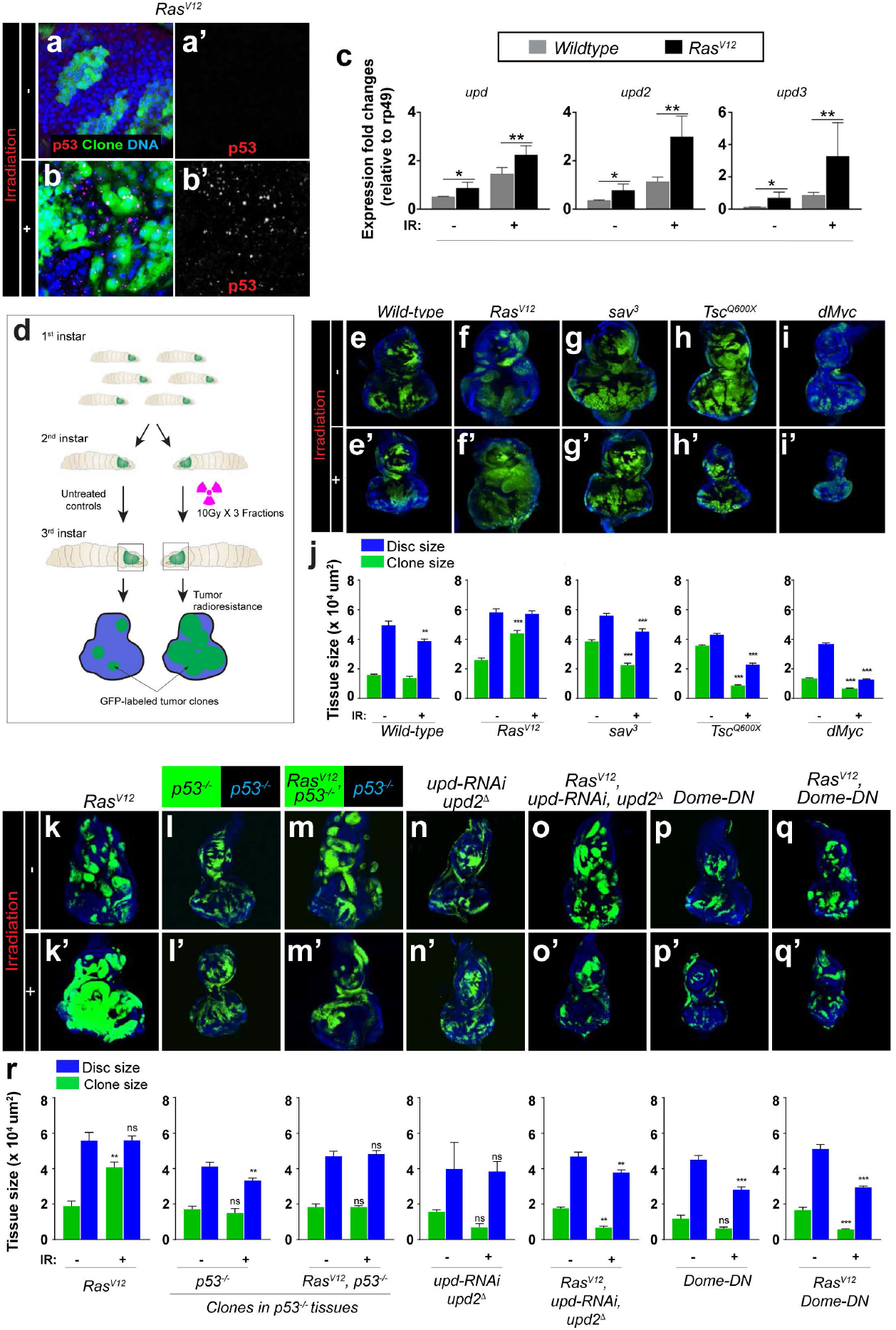
The Wild-type p53/oncogenic Ras nonautonomous STAT signal relay promotes the radioresistance of *Drosophila* Ras tumor tissues. (a-b’) Upregulated p53 within *Ras* clones after irradiation. GFP-labelled *Ras^V12^* clones were stained with anti-p53 antibody before irradiation (0 h) and 24 h after irradiation. Time was counted from the start of first faction of IR treatment. (c) Quantification by RT-qPCR of *upd*, *upd2*, and *upd3* expression in eye-antennal discs containing wild-type or *Ras^V12^* clones after 36 h of first fraction of IR treatment (IR+) or without IR treatment (IR-). Column bars represent the mean of fold changes for the expression level of indicated genes. Three independent experiments were carried out. Error bars denote SD. T-test significance levels are (*) *p*<0.05 and (**) *p*<0.01. (d) Diagram of setting *Drosophila* irradiation models. Larvae after egg-laying (48 h) were irradiated with three fractions of 10 Gy and allowed to recover to late third-instar larval stage. All eye discs were dissected at the late third-instar larval stage to evaluate the irradiation results by measuring the relative size between GFP-labeled clones and whole eye-antennal discs. (e-i’) GFP-labeled clones homozygous for *Ras^V12^* (f, f’), *sav^3^* (g, g’), *Tsc1^Q600X^* (h, h’), or expressing *dMyc* (i, i’) as well as wild-type controls (e, e’) were induced in the eye-antennal discs of larvae, irradiated at 48 h, and then discs collected discs on day 5. Upper panels (e-i) show the eye-antennal discs without irradiation treatment (IR-), and lower panels (e’-I’) show irradiated discs (IR+). (j) Quantification of relative eye disc size (blue) and GFP-clone size (green) treated with IR (IR+) or without IR (IR-). For each genotype, eye-antennal disc and GFP-clone were normalized to age-matched eye discs without IR. Column bars represent the mean size of samples (n, 5~10). Error bars denote SEM. t-test significance levels of are (*) *p*<0.05, (**) *p*<0.01, and (***) *p*<0.001. Numerical data are available in Figure 1 source data. (k-q’) GFP-labeled *Ras^V12^, p53^-/-^, Ras^V12^p53^-/-^, upd^RNAi^upd2*^Δ^, *Ras^V12^upd^RNAi^upd2*^Δ^, *Dome^DN^*, and *Ras^V12^ Dome^DN^* clones were induced in the eye-antennal discs and half were then irradiated at the second-instar larval stage. After 3 days of recovery, all eye discs at the late third-instar larval stage were dissected to evaluate the differences in response to irradiation. Upper panels (k-q) show the eye-antennal discs without irradiation treatment (IR-), and lower panels (k’-q’) show discs treated with irradiation (IR+). (r) Quantification of clones and eye discs treated with or without irradiation. Eye-antennal disc and GFP-clone areas were measured by ImageJ and normalized to the eye-antennal discs with the same genotype at the same age without IR. Column bars represent the mean size of samples (n=10). Blue columns represent the relative eye-antennal size with indicated genotypes; green columns represent the size of GFP-labeled tumors or non-tumor clones. Error bars denote SEM. T-test significance levels are (*) *p*<0.05, (**) *p*<0.01, and (***) *p*<0.001 (ns, not significant).

We sought to test the functional relevance of IR-induced STAT to *Ras^V12^* tumor radioresistance directly in Ras tissues. IR reduced the size of wild-type clones and the overall tissue size, as expected (Fig. 5e, e’, and j). In sharp contrast, IR increased *Ras^V12^* clone size and failed to reduce overall tissue size (Fig. 5f, f’, j, k, and k’). To determine whether this resistance is unique to oncogenic Ras signaling, we extended our analysis to other tumor signaling contexts: tissues containing clones of cells carrying homozygous null mutations in tumor suppressors (*Salvador/Sav* or *tuberous sclerosis/Tsc*) or tissues containing clones of cells overexpressing the oncogene *dMYC*. In all of these tissues, IR effectively reduced clone and overall tissue size (Fig. 5f-i’ and j). Thus, similar to human, *Drosophila* Ras tumors are uniquely radioresistant.

Next, we asked to what extent depletion of p53 or JAK/STAT cytokines sensitize *Ras^V12^* tissues to IR. We induced *Ras^V12^* clones in wild-type or *p53*^-/-^ null mutant discs, assessed whether the *p53*^-/-^ mutation sensitizes *Ras^V12^* tumor tissues to IR, and found that it does (Fig. 5k-m’, r).

Next, we simultaneously depleted Upd1 and Upd2 cytokines from *Ras^V12^* cells and asked whether this also sensitizes *Ras^V12^* tumor tissues to IR. Two independent approaches were used. First, we introduced an *upd2* null mutation into *Ras^V12^* cells coexpressing *Upd1-RNAi* (*Ras^V12^, upd^-/-^, Upd1-RNAi* clones). Second, we generated clones of cells coexpressing *Ras^V12^, upd-RNAi* and *upd2-RNAi*. Both approaches abolish the capacity of *Ras^V12^* tissues to grow following IR treatment, supporting a sensitization effect (Fig. 5k, k’, n-o’, and r). Similarly, inhibition of the JAK-STAT receptor *domeless* specifically in *Ras^V12^* cells via expression of a potent dominant-negative protein version (*domeless-DN*)^59^ sensitized *Ras^V12^* tissues to IR (Fig. 5k, k’, p-r).

Thus, in *Drosophila* and human cancer cells, p53 response to genomic instability cooperates with oncogenic Ras to induce JAK/STAT activity in surrounding cells. This nonautonomous effect stimulates tumor growth and promotes the rapid recurrence of oncogenic Ras tumors.

## Discussion

Oncogenic Ras mutations are associated with resistance to genotoxic therapies. The underlying resistance mechanism remains poorly understood. Tumor cells’ molecular responses to genotoxic stress have been investigated mainly in isolated cells, making it impossible to capture broader tissue level tumor biology.

Using the *Drosophila* eye tissue in a clonal genetic screen for *Ras^V12^* suppressors, we unexpectedly isolated the genotoxic mutation *ptip*^-/-^. Interestingly, in addition to blocking Ras tumor growth, *ptip*^-/-^ stimulates surrounding *Ras^V12^* tissue to overgrow, mirroring the resistance of *Ras^V12^* tumors to genotoxic therapies. This nonautonomous effect stems from a cooperation between oncogenic Ras signaling and *ptip*^-/-^, as oncogenic Ras or *ptip*^-/-^ mutant cells alone do not cause nonautonomous growth. Our mechanistic studies reveal that this cooperation is centered on p53.

P53 is broadly known as a tumor suppressor gene. The majority of *p53* mutations in human cancers are missense mutations that stabilize p53 protein, leading to elevated levels of mutant p53 protein in cancers. These gain-of-function mutations interfere with p53’s canonical tumor suppressor role while causing it to behave as an oncogene^60–63^. The accumulation of mutant p53 is associated with aggressive cancers^60–64^. Interestingly, overexpression of wild-type p53 is also observed in many cancers lacking *p53* mutations, including in lung cancers where oncogenic Ras mutations are common^65–69^. How wild-type p53 overexpression relates to oncogenic Ras cancers and their resistance to genotoxic therapies was unclear. Here, we show that genotoxic stress-activated p53 acts non-cell autonomously to promote the radioresistance of Ras tumor tissues.

The *ptip*^-/-^ mutation causes genomic instability in *Ras^V12^* cells, resulting in the upregulation of p53 protein. This stimulation of p53 is essential for the nonautonomous tissue overgrowth effect of *Ras^V12^ptip*^-/-^ tumor clones. RNAi depletion of p53 in *Ras^V12^ptip*^-/-^ clones abrogates the nonautonomous tissue overgrowth effect. In addition, direct overexpression of p53 in *Ras^V12^* clones is sufficient to trigger overgrowth of the surrounding tissues, mimicking the *ptip*^-/-^ mutation.

The nonautonomous tissue overgrowth effect is mediated by the p53 transcriptional program. Expression of a DNA binding defective p53 mutant (*p53^R155H^*)^70^ in *Ras^V12^ptip*^-/-^ cells blocks the nonautonomous tissue overgrowth effect. Congruent with this, the transcriptional program of wild-type p53 is modified in cancer-associated fibroblasts to promote cancer progression^71^. Also, under ectopic wild-type p53 conditions, oncogenic Ras modifies the p53 transcriptional program, leading to the senescence associated secretory phenotype (SASP)^72–74^.

JAK/STAT cytokines are the primary transcriptional targets relevant to the Ras/p53 cooperative effect on nonautonomous growth. JAK/STAT cytokines are transcriptionally upregulated in tissues showing nonautonomous growth (*Ras^V12^ptip*^-/-^ or *Ras^V12^p53^OE^*) compared to controls (*Ras^V12^*). Depleting Upd cytokines specifically in *Ras^V12^ptip*^-/-^ or *Ras^V12^p53^OE^* clones suppresses nonautonomous tissue overgrowth.

The cooperative effect of oncogenic Ras and p53 on paracrine STAT signaling may reflect an ability for Ras to rewire the p53 transcriptional program and/or to elevate the exocytosis of STAT cytokines above a required threshold. Consistent with the latter possibility, we identified and validated P53-binding sites near Upd genes. Interestingly, deletion of these sites using reporter assays significantly suppressed Upd expression but it did not restore Upd expression to basal levels (Supplementary Fig. 5), possibly because of cryptic P53 binding sites near Upd genes. p53 binds to non-canonical DNA sites to expand its transcriptional network^75^. In the future, it would be desirable to map these non-canonical p53 sites on Upd to better understand p53.

The nonautonomous Ras/P53-STAT signal relay allows Ras clones to resist the damaging effects of IR treatment in *Drosophila*, as p53 or STAT depletion sensitizes Ras tumor tissues to IR. Wild-type p53 is stimulated nonuniformly in irradiated Ras tissues. This heterogeneity could be due to stochastic variation or could reflect different cell-inherent capacities to successfully withstand genotoxic stress. Treatment-induced p53 heterogeneity within Ras tumor tissues would allow cells with extensive genomic insult (high p53 levels) to directly induce upregulation of JAK/STAT ligands, which instructs the nearby less-damaged (low p53) *Ras^V12^* cells to overproliferate and reestablish the tumor following treatment.

*In vitro* supernatant transfer and mouse xenograph experiments revealed a similar mechanism in human Ras cancer cells. Indeed, activation of STAT signaling is associated with resistance to genotoxic agents in human cancer cells^76–78^. We show that, compared to controls, media conditioned with irradiated or p53-overexpressing Ras cancer cells elevate STAT signaling and stimulate cell proliferation across genetically diverse cancer cells. We propose that the Ras-p53 nonautonomous STAT signal relay likely represents a tissue-level adaptive mechanism for selecting and expanding therapy-resistant tumor clones in the tumor microenvironment. This mechanism is reminiscent of oncogenic Ras paracrine activation of TGFa/amphiregulin signaling to establish resistance against EGFR blockade in colorectal cancers^79^.

In addition to highlighting an emerging role of p53 in cell-cell interactions, our findings provide a possible explanation for the paradoxical resistance of *Ras* cancers to genotoxic therapies despite having functional p53^80,81^. Thus, our data suggest that combining STAT inhibition with radiation therapy may improve clinical outcomes.

Developmental and regenerative signaling contexts may functionalize P53 in a similar fashion to maintain tissue homeostasis. Neighboring cells with different levels of wild-type p53 influence each other’s growth in *Drosophila* and mammalian tissues^82^.

## Methods

### Fly strains and generation of clones

Flies were raised on standard *Drosophila* media at 25 °C. Fluorescently labelled mitotic clones were produced in larval imaginal discs using the following strains: (1) *yw, eyFLP1; Act>y+>Gal4, UAS-GFP.S65T; FRT82B, Tub-Gal80;* (2) *yw*; *eyFLP5, Act>y+>Gal4, UAS-GFP; FRT82B, Tub-Gal80;* (3) *FRT42D; eyFLP6, Act>y+>Gal4, UAS-GFP;* (4) *yw,upd2*^Δ*3-62*^; *eyFLP5, Act>y+>Gal4, UAS-GFP; FRT82B, Tub-Gal80;* (5) *yw, eyFLP1; Act>y+>Gal4, UAS-GFP.S65T; FRT79E;* (6) *yw, eyFLP1; Act>y+>Gal4, UAS-GFP.S65T; Tub-Gal80, FRT79E;* and (7) *yw, eyFLP1; Act>y+>Gal4, UAS-GFP.S65T; Tub-Gal80, ptip^3804^, FRT79E*. Additional strains used were as follows: (1) *yw*; *FRT82B*; (2) *w*;*UAS-Ras^V12^*(II); (3) *w*; *UAS-Ras^V12^*(III); (4) *w*;; *FRT82B,Tsc1^Q600X^/TM6B;* (5) *yw*; *82B,sav^3^/TM3;* (6) *yw*; *UAS-dMyc; Sb/TM6B;* (7) *UAS-dp53/CyO* (gift from Nanami); (8) *yw; UAS-p53^R155H^/T(2;3)TSTL, CyO: TM6B, Tb* (Bloomington Stock Center, BL8419); (9) *yw; p53 ^5A-1-4^* (BL6815; (10) *UAS-p53-IR* (VDRC, v103001); (11) *yw;FRT42D,ubi-RFP.nls;* (12) *w,UAS-BskD^N^;* (13) *yw*; *UAS-Ras^V12^; FRT79E,ptip^3804^;* (14) *w*; *UAS-upd-IR(R1)* (III) (NIG5988R); (15) *yw,upd2*^Δ*3-62*^ (gift from M. Zeidler); (16) *yw,upd-lacZ* (gift from H. Sun); and (17) *w,UAS-dome*^Δ*cyt1.1*^ (gift from J. Hombria).

### *Drosophila* x-ray irradiation

Second-instar larvae (54±6 h after egg laying) were collected from a petri dish containing 3 ml food and irradiated with three fractions of 1000R at 6 h intervals in a Cabinet Faxitron X-Ray machine (TRX 2800). All larvae were allowed to recover at 25 °C. Eye discs were dissected at the late third-instar stage (day 5) based on developmental markers (spiracles and mouth hooks). For timing experiments, second-instar larvae were irradiated using three fractions of 1000R(10Gy), and dissections were timed from the end of treatment.

### Staining and imaging

Antibody staining was performed according to standard procedures for *Drosophila* imaginal discs. The following antibodies were used: mouse monoclonal anti-β gal (1:500, Sigma), mouse anti-dmp53 (1:50 DSHB), rabbit anti-Stat92E (1:1000; gift from S. Hou), rabbit anti p-JNK (1:100), and mouse anti-H2Av monoclonal antibody (1:200, DSHB #UNC93-5.2.1). Signals were detected by Alexa Fluor^®^ 568 Goat anti-mouse IgG (1:500, Life Technologies) or by Alexa Fluor^®^ 555 Goat anti-rabbit IgG (1:500, Life Technologies). Images were taken using a Leica SP8 confocal microscope. Measurements of tumor clone size within imaginal discs were performed from confocal pictures using Fiji imageJ software. Adult eyes were imaged with a Leica DFC 300FX camera in a Leica MZ FLIII fluorescence stereomicroscope.

### Real-time RT-PCR

Total RNA from wild-type and tumor imaginal discs (n≥30 pairs) were extracted using Trizol. cDNAs were synthesized from 500 ng total RNA with the iScript cDNA Synthesis Kit (Bio-Rad). cDNAs were subjected to real-time PCR with the SYBR Green Fast Kit (Applied Biosystems), according to the manufacturer’s instructions. The expression level of genes for each sample was calculated by comparing to the internal control, rp49. The relative fold change of each gene was normalized to the expression level of the same gene in the eye disc bearing *Ras^V12^*.

Three experiments for each condition were averaged. The following primers were used for qRT-PCR:

Upd: 5’-TCCACACGCACAACTACAAGTTC-3’; 5’-CCAGCGCTTTAGGGCAATC-3’
upd2: 5’-AGTGCGGTGAAGCTAAAGACTTG-3’;5’-GCCCGTCCCAGATATGAGAA-3’
upd3: 5’-TGCCCCGTCTGAATCTCACT-3’; 5’-GTGAAGGCGCCCACGTAA-3’
rp49: 5’-GGCCCAAGATCGTGAAGAAG-3’; 5’-ATTTGTGCGACAGCTTAGCATATC-3’

### Luciferase assay

The XmaI-XbaI fragment, including the predicted p53 binding sites, from upstream of Upd2 was separated from BAC DNA (BACR32F24) by enzyme digestion and then ligated into the pGL3-promotor to get the pGL3-Upd2-luc reporter construct (Upd2-p53BDS). The upstream fragment of Upd3 was separated by XmaI-SpeI enzyme digestion from BAC DNA and then ligated into the pGL3-promoter plasmid to form the pGL3-upd3-luc (Upd3-p53BDS). The predicted p53 binding sites were deleted from the reporter constructs to generate pGL3-upd2^Δ^-luc (Upd2-p53BDS^Δ^) and pGL3-upd3^Δ^-luc (Upd3-p53BDS^Δ^). 1.6×10^5^ Schneider S2R+ cells per well were plated in a 24-well plate one day before transfection. S2R+ cells were transfected with pGL3-Upd2-luc and pGL3-Upd3-luc reporter constructs and the p53 binding site deletion constructs separately along with metallothionein promoter-dp53 constructs (MT-p53) (from DGRC, FMO05476) and Renilla reporter (pRL-TK Vector) by using the X-treme GENE HP DNA Transfection Reagent (Roche). After 48 h of transfections, copper sulphate was added to the medium in a final concentration 500 μM to induce dp53 expression. After the inducing expression of dp53 for 24 h, a dual luciferase assay was performed. Relative expression levels were calculated based on the Renilla reporter as an internal control. Experiments were done in triplicates.

### Cell lines and cell culture

H441, H358, H460, and A549 were authenticated at the Cell and Immunology Core facility at the University of Missouri. MCF-10A and MCF-10A^Onc HRAS^ cells were a gift from G. Monogarov (DKFZ, Heidelberg, Germany). Cells were cultured following ATCC recommendation in DMEM media containing high glucose and L-glutamine (Gibco#11875-093) or in RPMI-1640 media for H441 cells. A549 cells were cultured in Dulbecco’s Modified Eagle Medium with L-glutamine and high glucose (DMEM#11965-092) supplemented with 10% FBS. MCF-10A cells were cultured in DMEM-F12 (11320-033) with 5% horse serum (Gibco, 16050-122), hydrocortisone 0.5 μg/ml, cholera toxin 100 ng/ml, insulin 10 μg/ml, and EGF 20 ng/ml. Cells were incubated in a humidified incubator with 5% CO2 at 37 °C.

### Cell counting

TrypLE (Gibco#12604-021) was used for detaching cells. Cells were counted using Trypan Blue as a dead cell excluder (Cat #T8154, Sigma USA).

### Transfection

The p53-GFP vector was purchased from addgene (#1209). DharmaFECT (Dharmacon# T2002-02) was used for transfecting the p53-GFP vector into the human cell lines, following the manufacturer’s recommendation.

### Ganetespib cytotoxicity

MTT assays on cancer cells treated with increasing concentrations of Ganetespib (0.1 to 100 nM) determined the half maximal inhibitory concentration (IC50) at 25 nM. Cells were seeded in 96-well plates at a density of 1×10^5^ cells per well and treated with Ganetespib (Biosciences, #A11402,) for 48 h. A 100 μl solution of MTT (100 μg/ml) was added to each well. Cell viability was measured with a spectrophotometer at 570 nm.

### Immunoblotting

For the immunoblotting analysis, cells were lysed in a buffer (20 mM Tris-HCl pH-7.5, 150 mM NaCl, 1 mM Na2EDTA, 1 mM EGTA, 1% Triton, 2.5 mM sodium pyrophosphate, 1mM b-glycerophosphate, 1mM Na3VO4, 1 μg/ml leupeptin) supplemented with protease and a phosphatase inhibitor cocktail (Cell Signaling #9803S). Absolute protein concentration was determined and normalized using a BSA standard curve on Nanodrop (Thermo Fisher Scientific). Proteins were electrophoresed on SDS-PAGE using 4-15% Mini-PROTEAN^®^ TGX^™^ precast gel (Biorad#456-1085) and transferred onto a nitrocellulose membrane using Trans-Blot^®^ Turbo^™^ Transfer Pack (Biorad#170-4159). The nitrocellulose membrane was blocked in a buffer (5% milk in Tris-buffered saline with containing 0.01% Tween20). Membranes were probed overnight at 4 °C with anti-pSTAT-3 (Tyr 705) (1:1000, Cell Signaling #9145) or anti-STAT (1:1000, Cell Signaling #30835) or anti-GAPDH (1:1000, Cell signaling #5174S). Secondary horseradish peroxidase (HRP) antibodies were obtained from Invitrogen (1:5000, Invitrogen#31460). An ECL Chemiluminescence Kit (Thermo Fisher Scientific #32106) and the ChemiDoc Imaging System (Bio-Rad) were used to detect protein bands.

### Human cancer cells irradiation

For x-irradiation conditioned media experiments, cells were cultured in fresh media prior to irradiation (8 Gy, 280 cGy/min exposer) using an X-RAD 320 Biological Irradiator. Media were replaced immediately following irradiation, and the irradiated cells were cultured for 24 h to generate conditioned media. Supernatant from the irradiated cells was collected after centrifugation at 500 xg for 2 mins to remove cellular debris.

### Flow cytometry analysis

GFP cell sorting was performed on the MoFloXDP (Beckman Coulter), and the regular flow cytometer analysis was performed using a CyAn ADP (Beckman Coulter). Cell proliferation flow cytometry assays were performed using the 5-ethynyl-2’deoxyuridine (EdU) assay kit (Invitrogen #C10634). Cells were incubated with EdU for 24 h and labeled. The Click-iT reaction was performed using the Click-iT EdU assay kit with Alexa Fluor 647 fluorophore, according to the manufacturer’s instructions. The propidium iodide (PI) solution was at 10 μg/ml in PBS containing 1% BSA. PI was detected at 488/636 nm (excitation/emission) and EdU-Alexa647 at 633/660 nm.

### Mouse xenograft experiments

Thirteen-week-old athymic nude mice (homozygous *Foxn1^nu^*) were purchased from the Jackson Laboratory for xenograph experiments. Experiments were conducted in compliance with the National Institute of Health’s guide for the care and use of animals. All animals were housed under pathogen-free conditions on a 12/12 h light-dark cycle. A549 lung cancer cells were collected. The A549 cells were grown in Dulbecco’s Modified Eagle Medium with L-glutamine and high glucose supplemented with 10% FBS. Cells were collected in pharmaceutical grade PBS, counted, and resuspended in pharmaceutical grade PBS at 1×10^6^/100 uL. All the left flanks were inoculated with 1×10^6^ A549 cells. Right flanks were inoculated with a 50/50 mixture of A549 and irradiated A549 (0.5×10^6^ +0.5×10^6^ cells) in a total of 100 uL. Tumor measurements were taken weekly using a digital caliper. The STAT inhibitor Ruxolitinib (Sigma Millipore #ADV390218177) was administered (10 mg/kg body weight) through oral gavage once a day for a period of three weeks.

### Statistical analysis

All *in vitro* experiments were performed in three biological replicates for reproducibility of data. Each data point with standard deviations represents at least three replicates. Student’s *t-test* was used to determine statistical significance of differences between groups.

## Author Contributions

YD, TK, JL, PG, TX, and CYC conceived the study. YD and TK developed the fly irradiation models. LL assisted with qRT-PCR and fly stock maintenance. XM assisted with immunofluorescence staining experiments. GP performed the tissue culture and mouse xenograph experiments. YD and CYC wrote the manuscript.

## Acknowledgements

We thank M. Oren, G. Monogarov, J. Contessa, B. Dunn, S. Ding, F. Qian, H. Chang, and D. Manry for helpful discussions. We are grateful to M. Oren and G. Monogarov for providing MCF-10A cells. We also thank N. Senoo-Matsuda, H. Zeidler, J. Sun, S. Hombria, S. Hou, the Bloomington Stock Center, the Vienna *Drosophila* RNAi Center, and the National Institute of Genetics for fly strains and antibodies. TK was supported by an RSNA research grant. This work was supported in part by the Chinese National Natural Science Funds for Young Scholars (No. 31200687) to YD, a NIH/NCI grant to TX and the Howard Hughes Medical Institute, and startup funds to CYC from the University of Missouri.

---

**Supplementary Figure 1.**
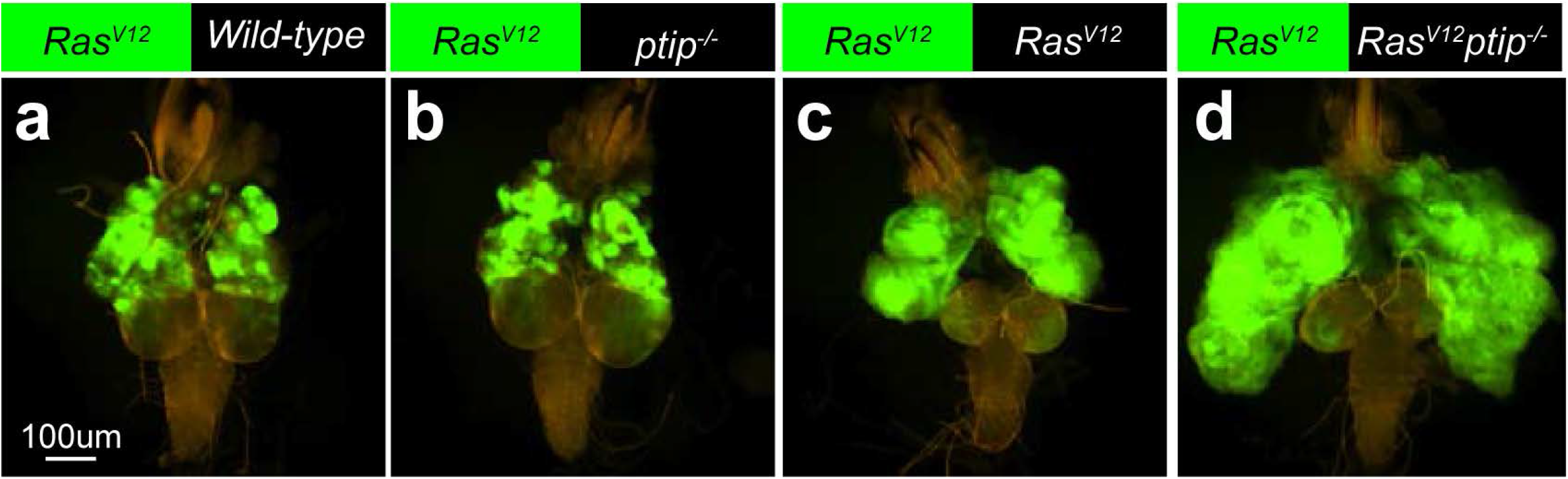
Oncogenic Ras and the *ptip*^-/-^ mutation cooperate to accelerate the growth of nearby Ras clones. Images showing the growth of mosaic eye imaginal tissue from dissected brain complexes. Eye imaginal tissues contain GFP-positive *Ras^V12^* clones juxtaposed against wild-type (a) or *ptip*^-/-^ (b) or *Ras^V12^* (c) or *Ras^V12^*ptip^-/-^ (d) mutant clones.

**Supplementary Figure 2.**
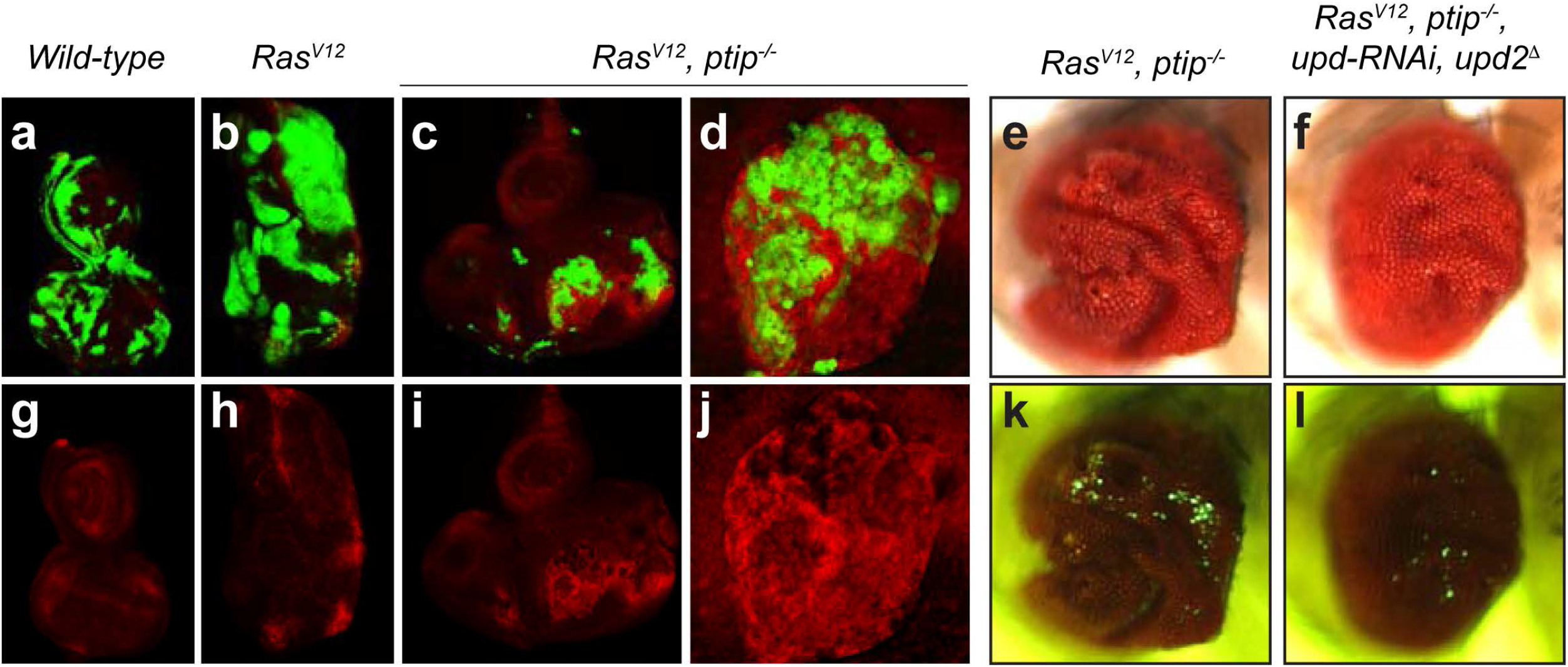
Oncogenic Ras and *ptip*^-/-^ stimulate nonautonomous tissue overgrowth via paracrine activation of STAT signaling. (a-d) Representative images of dissected eye imaginal tissues containing GFP-labeled clones of wild-type (a) or *Ras^V12^* (b) or *Ras^V12^ptip*^-/-^ (c-d) mutant cells stained with an antibody that detects STAT92E (red). Corresponding individual STAT92E channel images are shown in bottom panels (g-j). (e-l) Matched light and fluorescence images of adult eyes containing *Ras^V12^ptip*^-/-^ mutant clones in the absence (e, k) or presence (f, l) of *upd-RNAi* and *upd*2 deletion (upd2^□^)

**Supplementary Figure 3.**
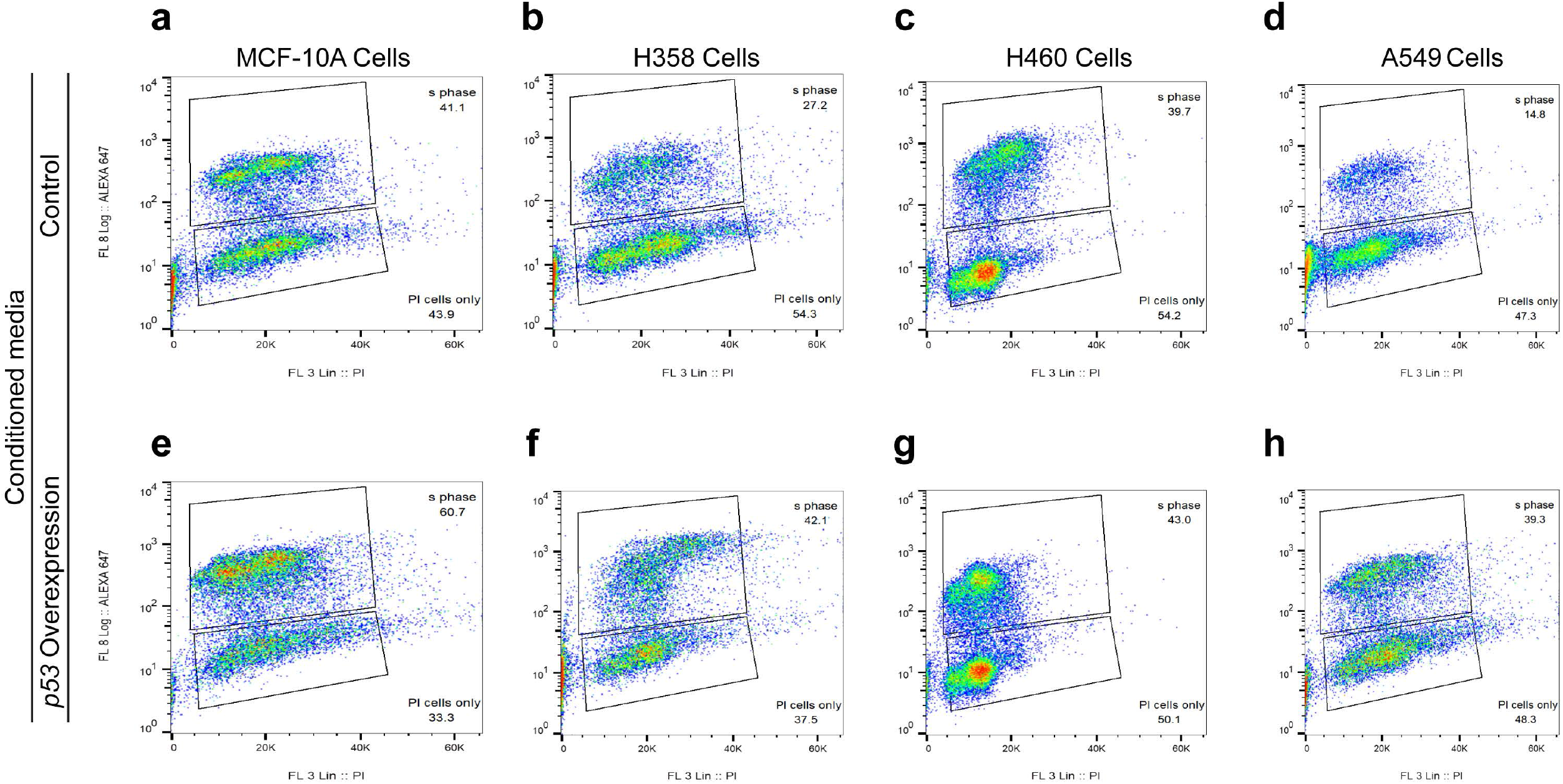
Media conditioned with oncogenic Ras cells overexpressing *p53* elevate the cell proliferation potential of human cancer cells. (a-h) Flow cytometry plots from experiments using the indicated cancer cells. Cancer cells were co-labeled with EDU-647 and propidium iodide (PI) to determine the proportion of proliferating cells and dying cells, respectively. Top row (a-d) shows the proportion of proliferating and dying cells when cells are cultured in control media (conditioned with unmodified matching cells). Bottom row shows the proportion of proliferating and dying cells when cells are cultured in media conditioned with matching *p53*-overexpressing cells.

**Supplementary Figure 4.**
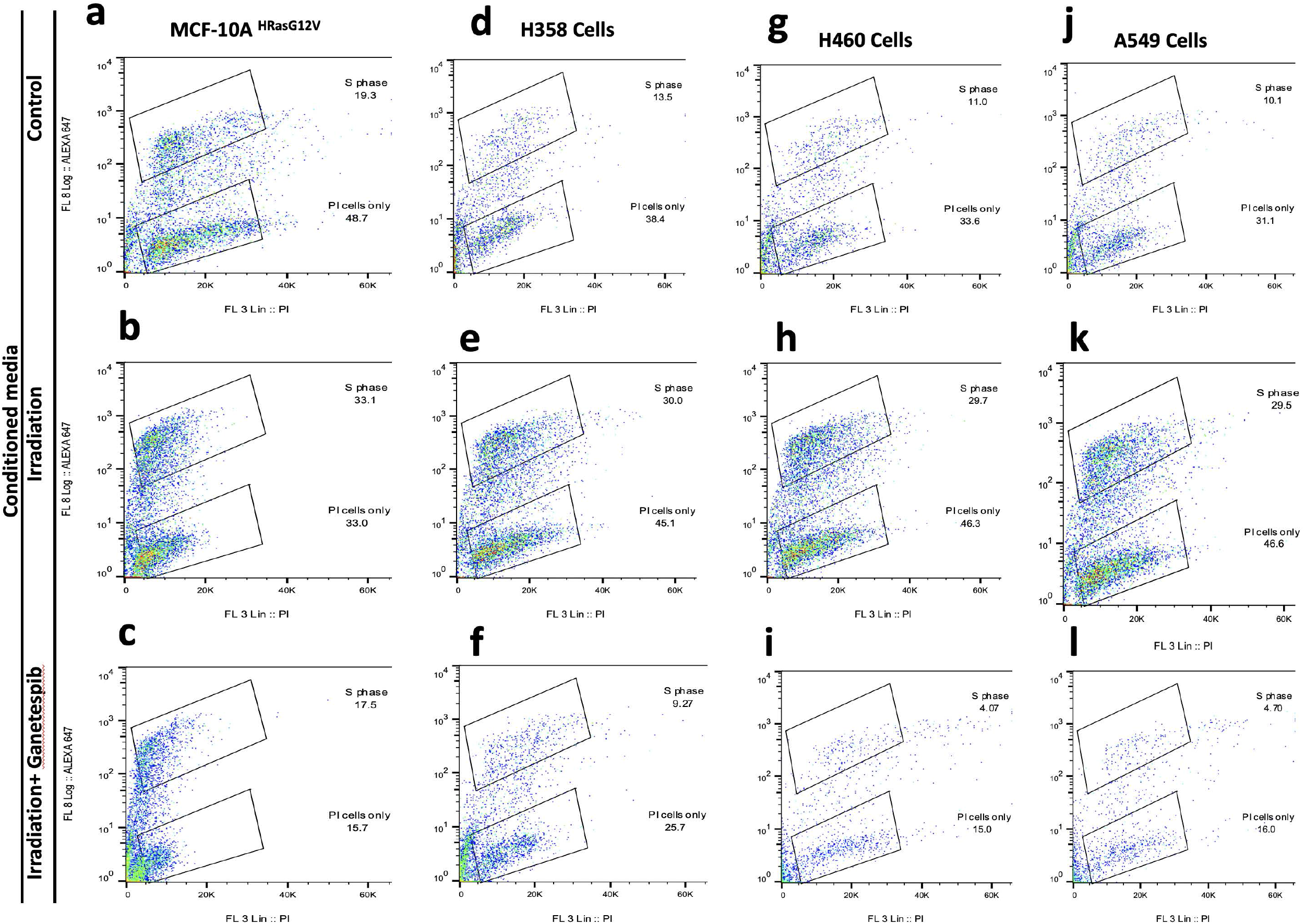
Media conditioned with irradiated cells expressing oncogenic Ras elevate the cell proliferation potential of human cancer cells. (a-l) Flow cytometry plots from experiments using the indicated cancer cells. Cancer cells were co-labeled with EDU-647 and propidium iodide (PI) to determine the proportion of proliferating cells and dying cells, respectively. The top row (a, d, g, and j) shows the proportion of proliferating and dying cells when cells are cultured in media conditioned with unmodified matching cells (baseline controls). The middle row (b, e, h, and k) shows the proportion of proliferating and dying cells when cells are cultured in media conditioned with the matching irradiated cells. The bottom row (c, f, i, and l) shows the proportion of proliferating and dying cells when cells are cultured in media conditioned with matched irradiated cells in the presence of Ganetespib.

**Supplementary Figure 5.**
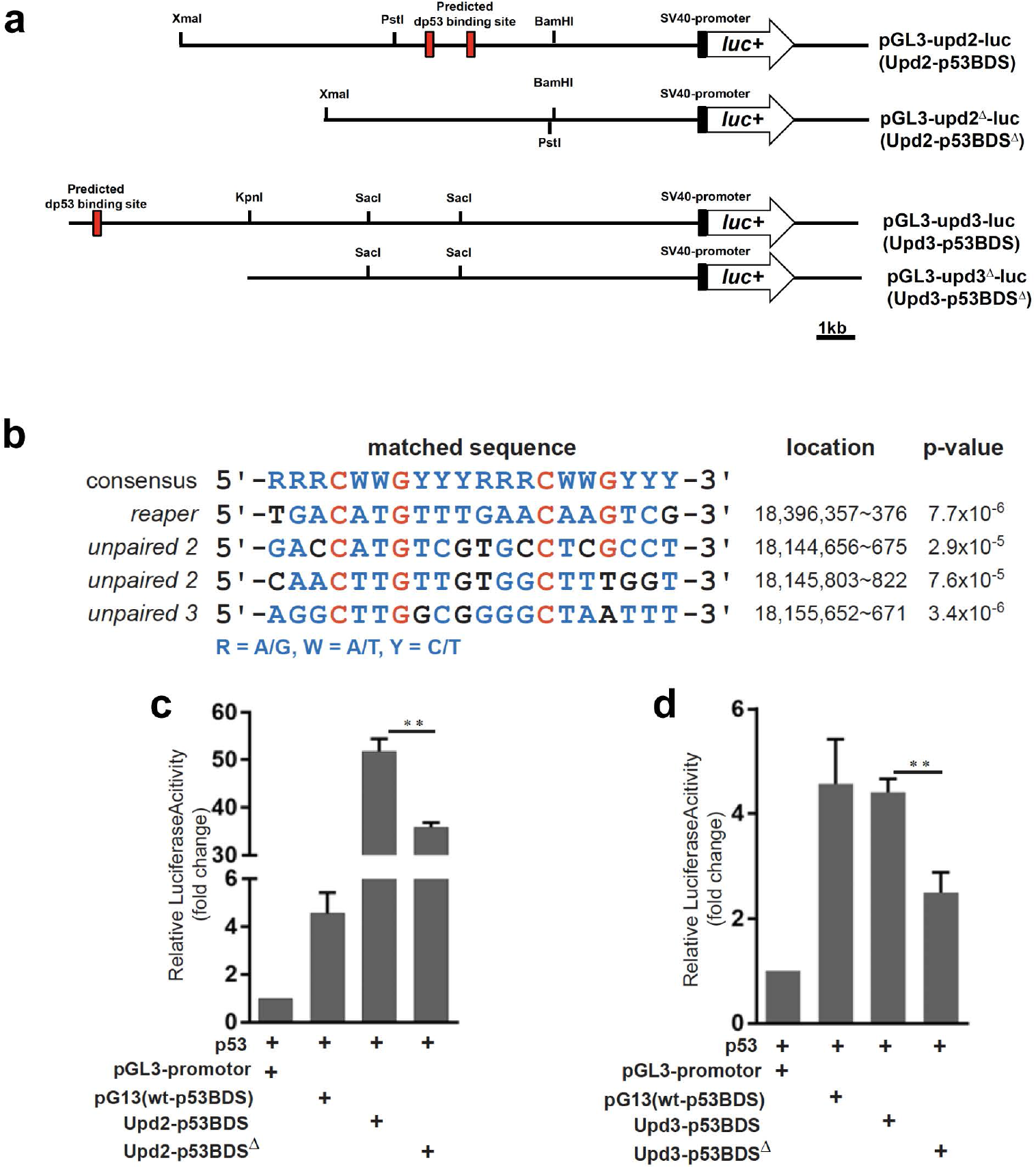
*In silico* identification of functional p53 binding sequences upstream of unpaired genes. (a) Graphical representation of the strategy used to develop the *upd* promoter activity reporter assay. Putative p53 binding sites (p53BDS) from the upstream regions of *upd2* and *upd3* genes are denoted with red boxes. (b) Genomic sequences showing *reaper* consensus p53BDS matched to unpaired 2 and unpaired 3 upstream sequences. Note that the p53BDS from upstream of reaper gene is a well-defined p53 sequence. Corresponding locations and p-values are shown to the right. (c, d) Luciferase activity in S2R^+^ cells after copper sulfate induction of p53 expression in the absence or presence of Upd2-p53BDS, Upd3-p53BDS, as well as p53BDS-deleted control (Upd2-p53BDS^□^, Upd3-p53BDS^□^) sequences. T-test significance levels are (*)*p<0.05* and (**) *p<0.01*.

## Notes

### Competing Interest Statement

The authors have declared no competing interest.

## References

1. Cox, A. D., Fesik, S. W., Kimmelman, A. C., Luo, J. & Der, C. J. Drugging the undruggable RAS: Mission Possible ? Nature Publishing Group 13, 828–851 (2014).

2. Hanahan, D. et al. The Hallmarks of Cancer. Cell 100, 57–70 (2014).

3. Zerdoumi, Y. et al. A new genotoxicity assay based on p53 target gene induction. Mutat Res Genet Toxicol Environ Mutagen 789–790, 28–35 (2015).

4. Bernhard, E. J. et al. Direct Evidence for the Contribution of Activated N-ras and K-ras Oncogenes to Increased Intrinsic Radiation Resistance in Human Tumor Cell Lines. Cancer research 60, 6597–6600 (2000).

5. Gupta, A. K. et al. The Ras Radiation Resistance Pathway. 4278–4282 (2001).

6. Miller, A. C., Kariko, K., Myers, C. E., Clark, E. P. & Samid, D. Increased radioresistance of EJras-transformed human osteosarcoma cells and its modulation by lovastatin, an inhibitor of p21ras isoprenylation. Int J Cancer 53, 302–307 (1993).

7. Ling, C. C. & Endlich, B. Radioresistance induced by oncogenic transformation. Radiat Res 120, 267–279 (1989).

8. Garrido, P. et al. Treating KRAS-mutant NSCLC: latest evidence and clinical consequences. Ther Adv Med Oncol 9, 589–597 (2017).

9. Mak, R. H. et al. Outcomes by tumor histology and kras mutation status after lung stereotactic body radiation therapy for early-stage non-small-cell lung cancer. Clinical Lung Cancer 16, 24–32 (2015).

10. Affolter, A. et al. Increased radioresistance via G12S K-Ras by compensatory upregulation of MAPK and PI3K pathways in epithelial cancer. Head Neck 35, 220–228 (2013).

11. Minjgee, M., Toulany, M., Kehlbach, R., Giehl, K. & Rodemann, H. P. K-RAS(V12) induces autocrine production of EGFR ligands and mediates radioresistance through EGFR-dependent Akt signaling and activation of DNA-PKcs. Int J Radiat Oncol Biol Phys 81, 1506–1514 (2011).

12. Caron, R. W. et al. H-RAS V12-induced radioresistance in HCT116 colon carcinoma cells is heregulin dependent. Mol Cancer Ther 4, 243–255 (2005).

13. Ziv, E. et al. Lung Adenocarcinoma: Predictive Value of KRAS Mutation Status in Assessing Local Recurrence in Patients Undergoing Image-guided Ablation. Radiology 282, 251–258 (2017).

14. Bussink, J., van der Kogel, A. J. & Kaanders, J. H. Activation of the PI3-K/AKT pathway and implications for radioresistance mechanisms in head and neck cancer. Lancet Oncol 9, 288–296 (2008).

15. Bernhard, E. J. et al. Inhibiting ras prenylation increases the radiosensitivity of human tumor cell lines with activating mutations of ras oncogenes. Cancer Research 58, 1754–1761 (1998).

16. Gupta, A. K. et al. The Ras radiation resistance pathway. Cancer Res 61, 4278–4282 (2001).

17. Rait, A. et al. 3’-End conjugates of minimally phosphorothioate-protected oligonucleotides with 1-O-hexadecylglycerol: Synthesis and anti-ras activity in radiation-resistant cells. Bioconjugate Chemistry 11, 153–160 (2000).

18. Wang, M. et al. Radiation Resistance in KRAS-Mutated Lung Cancer Is Enabled by Stem-like Properties Mediated by an Osteopontin – EGFR Pathway. 77, 2018–2029 (2017).

19. Ramachandran, S., Ramadas, K., Hariharan, R., Rejnish Kumar, R. & Radhakrishna Pillai, M. Single nucleotide polymorphisms of DNA repair genes XRCC1 and XPD and its molecular mapping in Indian oral cancer. Oral Oncol 42, 350–362 (2006).

20. Bao, Y. et al. XRCC1 gene polymorphisms and the risk of differentiated thyroid carcinoma (DTC): a meta-analysis of case-control studies. PLoS One 8, e64851 (2013).

21. Zhai, X. M. et al. Significance of XRCC1 Codon399 polymorphisms in Chinese patients with locally advanced nasopharyngeal carcinoma treated with radiation therapy. Asia Pac J Clin Oncol 12, e125–32 (2016).

22. Yu, J. J. et al. Comparison of two human ovarian carcinoma cell lines (A2780/CP70 and MCAS) that are equally resistant to platinum, but differ at codon 118 of the ERCC1 gene. Int J Oncol 16, 555–560 (2000).

23. Park, D. J. et al. A Xeroderma pigmentosum group D gene polymorphism predicts clinical outcome to platinum-based chemotherapy in patients with advanced colorectal cancer. Cancer Res 61, 8654–8658 (2001).

24. Saranath, D. et al. High frequency mutation in codons 12 and 61 of H-ras oncogene in chewing tobacco-related human oral carcinoma in India. Br J Cancer 63, 573–578 (1991).

25. Anderson, J. A., Irish, J. C., McLachlin, C. M. & Ngan, B. Y. H-ras oncogene mutation and human papillomavirus infection in oral carcinomas. Arch Otolaryngol Head Neck Surg 120, 755–760 (1994).

26. Mladenov, E., Li, F., Zhang, L., Klammer, H. & Iliakis, G. Intercellular communication of DNA damage and oxidative status underpin bystander effects. Int J Radiat Biol 94, 719–726 (2018).

27. Chi, C. et al. Disruption of lysosome function promotes tumor growth and metastasis in Drosophila. J Biol Chem 285, 21817–21823 (2010).

28. Pagliarini, R. a & Xu, T. A genetic screen in Drosophila for metastatic behavior. Science (New York, NY) 302, 1227–1231 (2003).

29. Munoz, I. M., Jowsey, P. A., Toth, R. & Rouse, J. Phospho-epitope binding by the BRCT domains of hPTIP controls multiple aspects of the cellular response to DNA damage. Nucleic Acids Res 35, 5312–5322 (2007).

30. Jowsey, P. a, Doherty, A. J. & Rouse, J. Human PTIP facilitates ATM-mediated activation of p53 and promotes cellular resistance to ionizing radiation. The Journal of biological chemistry 279, 55562–9 (2004).

31. Gong, Z., Cho, Y. W., Kim, J. E., Ge, K. & Chen, J. Accumulation of pax2 transactivation domain interaction protein (PTIP) at sites of DNA breaks via RNF8-dependent pathway is required for cell survival after DNA damage. Journal of Biological Chemistry 284, 7284–7293 (2009).

32. Lee, J. et al. A tumor suppressive coactivator complex of p53 containing ASC-2 and histone H3-lysine-4 methyltransferase MLL3 or its paralogue MLL4. Proceedings of the National Academy of Sciences of the United States of America 106, 8513–8518 (2009).

33. Kastenhuber, E. R. & Lowe, S. W. Putting p53 in Context. Cell 170, 1062–1078 (2017).

34. Bieging, K. T., Mello, S. S. & Attardi, L. D. Unravelling mechanisms of p53-mediated tumour suppression. Nature reviews Cancer 14, 359–70 (2014).

35. Levine, A. J. & Oren, M. The first 30 years of p53: growing ever more complex. Nat Rev Cancer 9, 749–758 (2009).

36. Vousden, K. H. & Prives, C. Blinded by the Light: The Growing Complexity of p53. Cell 137, 413–31 (2009).

37. Chabu, C. & Xu, T. Oncogenic Ras stimulates Eiger/TNF exocytosis to promote growth. Development (Cambridge, England) 141, 4729–39 (2014).

38. Karim, F. D. & Rubin, G. M. Ectopic expression of activated Ras1 induces hyperplastic growth and increased cell death in Drosophila imaginal tissues. Development 125, 1–9 (1998).

39. Halfar, K., Rommel, C., Stocker, H. & Hafen, E. Ras controls growth, survival and differentiation in the Drosophila eye by different thresholds of MAP kinase activity. Development 128, 1687–1696 (2001).

40. Lechner, M. S., Levitan, I. & Dressler, G. R. PTIP, a novel BRCT domain-containing protein interacts with Pax2 and is associated with active chromatin. Nucleic acids research 28, 2741–2751 (2000).

41. Bouchard, M., Souabni, A., Mandler, M., Neubuser, A. & Busslinger, M. Nephric lineage specification by Pax2 and Pax8. Genes Dev 16, 2958–2970 (2002).

42. Torres, M., Gomez-Pardo, E., Dressler, G. R. & Gruss, P. Pax-2 controls multiple steps of urogenital development. Development 121, 4057–4065 (1995).

43. Cho, E. A., Prindle, M. J. & Dressler, G. R. BRCT domain-containing protein PTIP is essential for progression through mitosis. Molecular and cellular biology 23, 1666–1673 (2003).

44. Madigan, J. P., Chotkowski, H. L. & Glaser, R. L. DNA double-strand break-induced phosphorylation of Drosophila histone variant H2Av helps prevent radiation-induced apoptosis. Nucleic acids research 30, 3698–3705 (2002).

45. Lake, C. M., Holsclaw, J. K., Bellendir, S. P., Sekelsky, J. & Hawley, R. S. The development of a monoclonal antibody recognizing the Drosophila melanogaster phosphorylated histone H2A variant (γ-H2AV). G3 (Bethesda, Md) 3, 1539–43 (2013).

46. Vogelstein, B., Lane, D. & Levine, A. J. Surfing the p53 network. Nature 408, 307–310 (2000).

47. Karin, M. & Gallagher, E. From JNK to pay dirt: jun kinases, their biochemistry, physiology and clinical importance. IUBMB Life 57, 283–295 (2005).

48. Wagner, E. F. & Nebreda, A. R. Signal integration by JNK and p38 MAPK pathways in cancer development. Nat Rev Cancer 9, 537–549 (2009).

49. Brooks, C. L. & Gu, W. p53 ubiquitination: Mdm2 and beyond. Mol Cell 21, 307–315 (2006).

50. Wu, M., Pastor-Pareja, J. C. & Xu, T. Interaction between Ras(V12) and scribbled clones induces tumour growth and invasion. Nature 463, 545–548 (2010).

51. Ohsawa, S. et al. Mitochondrial defect drives non-autonomous tumour progression through Hippo signalling in Drosophila. Nature 490, 547–551 (2012).

52. Sun, Y. H. et al. white as a reporter gene to detect transcriptional silencers specifying position-specific gene expression during Drosophila melanogaster eye development. Genetics 141, 1075–1086 (1995).

53. Tsai, Y. C. & Sun, Y. H. Long-range effect of Upd, a ligand for Jak/STAT pathway, on cell cycle in Drosophila eye development. Genesis 39, 141–153 (2004).

54. Blanco, R. et al. A gene-alteration profile of human lung cancer cell lines. Hum Mutat 30, 1199–1206 (2009).

55. Proia, D. A. et al. Multifaceted intervention by the Hsp90 inhibitor ganetespib (STA-9090) in cancer cells with activated JAK/STAT signaling. PLoS One 6, e18552 (2011).

56. Mesa RA, Y. U. et al. Ruxolitinib. Nature reviews Drug Discovery 11, 103–104 (2012).

57. Tavallai M, Booth L, Roberts JL, Poklepovic, D. P. Rationally Repurposing Ruxolitinib (Jakafi (®)) as a Solid Tumor Therapeutic. Front Oncol 13, 142 (2016).

58. Bromberg, J. F. Activation of STAT proteins and growth control. Bioessays 23, 161–169 (2001).

59. Bromberg, J. F., Horvath, C. M., Wen, Z., Schreiber, R. D. & Darnell Jr., J. E. Transcriptionally active Stat1 is required for the antiproliferative effects of both interferon alpha and interferon gamma. Proc Natl Acad Sci U S A 93, 7673–7678 (1996).

60. La Fortezza, M. et al. JAK/STAT signalling mediates cell survival in response to tissue stress. Development 143, 2907–2919 (2016).

61. Thames, H. D., Bentzen, S. M., Turesson, I., Overgaard, M. & Van den Bogaert, W. Time-dose factors in radiotherapy: a review of the human data. Radiotherapy and Oncology 19, 219–235 (1990).

62. Barnett, G. C. et al. Normal tissue reactions to radiotherapy: towards tailoring treatment dose by genotype. Nature reviews Cancer 9, 134–42 (2009).

63. Brown, S., Hu, N. & Hombría, J. C. G. Identification of the first invertebrate interleukin JAK/STAT receptor, the Drosophila gene domeless. Current Biology 11, 1700–1705 (2001).

64. Oren, M. & Rotter, V. Mutant p53 gain-of-function in cancer. Cold Spring Harbor perspectives in biology 2, a001107 (2010).

65. Freed-Pastor, W. A. & Prives, C. Mutant p53: one name, many proteins. Genes Dev 26, 1268–1286 (2012).

66. Muller, P. A. & Vousden, K. H. p53 mutations in cancer. Nat Cell Biol 15, 2–8 (2013).

67. Brosh, R. & Rotter, V. When mutants gain new powers: news from the mutant p53 field. Nat Rev Cancer 9, 701–713 (2009).

68. Terzian, T. et al. The inherent instability of mutant p53 is alleviated by Mdm2 or p16INK4a loss. Genes Dev 22, 1337–1344 (2008).

69. Wang, Y. C. et al. Wild-type p53 overexpression and its correlation with MDM2 and p14ARF alterations: an alternative pathway to non-small-cell lung cancer. J Clin Oncol 23, 154–164 (2005).

70. Pardo, F. S. et al. Mutant, wild type, or overall p53 expression: freedom from clinical progression in tumours of astrocytic lineage. Br J Cancer 91, 1678–1686 (2004).

71. Houben, R. et al. High-level expression of wild-type p53 in melanoma cells is frequently associated with inactivity in p53 reporter gene assays. PLoS One 6, e22096 (2011).

72. Prior, I. A., Lewis, P. D. & Mattos, C. A comprehensive survey of Ras mutations in cancer. Cancer Res 72, 2457–2467 (2012).

73. Cancer Genome Atlas Research, N. Comprehensive molecular profiling of lung adenocarcinoma. Nature 511, 543–550 (2014).

74. Tsuchida, N., Murugan, A. K. & Grieco, M. Kirsten Ras* oncogene: significance of its discovery in human cancer research. Oncotarget 7, 46717–46733 (2016).

75. Kurtkaya-Yapicier, O., Scheithauer, B. W., Hebrink, D. & James, C. D. p53 in nonneoplastic central nervous system lesions: an immunohistochemical and genetic sequencing study. Neurosurgery 51, 1245–1246 (2002).

76. Ollmann, M. et al. Drosophila p53 is a structural and functional homolog of the tumor suppressor p53. Cell 101, 91–101 (2000).

77. Arandkar, S. et al. Altered p53 functionality in cancer-associated fibroblasts contributes to their cancer-supporting features. Proc Natl Acad Sci U S A 115, 6410–6415 (2018).

78. Coppé, J.-P. et al. Senescence-associated secretory phenotypes reveal cell-nonautonomous functions of oncogenic RAS and the p53 tumor suppressor. PLoS biology 6, 2853–68 (2008).

79. Ferbeyre, G. et al. Oncogenic ras and p53 Cooperate To Induce Cellular Senescence Oncogenic ras and p53 Cooperate To Induce Cellular Senescence. (2002). doi:10.1128/MCB.22.10.3497

80. Nakamura, M., Ohsawa, S. & Igaki, T. Mitochondrial defects trigger proliferation of neighbouring cells via a senescence-associated secretory phenotype in Drosophila. Nature Communications 5, 5264 (2014).

81. Menendez, D., Inga, A. & Resnick, M. A. The expanding universe of p53 targets. Nat Rev Cancer 9, 724–737 (2009).

82. Spitzner, M. et al. STAT3: A novel molecular mediator of resistance to chemoradiotherapy. Cancers 6, 1986–2011 (2014).

83. Wu, C., Chen, M., Chen, W. & Hsieh, C. The role of IL-6 in the radiation response of prostate cancer. Radiation Oncology 8, 1 (2013).

84. Hu Y, Hong Y, Xu Y, Liu P, Guo DH, C. Y. Inhibition of the JAK/STAT pathway with ruxolitinib overcomes cisplatin resistance in non-small-cell lung cancer NSCLC. Apoptosis 19, 1627–36 (2014).

85. Hobor, S. et al. TGFalpha and amphiregulin paracrine network promotes resistance to EGFR blockade in colorectal cancer cells. Clin Cancer Res 20, 6429–6438 (2014).

86. Hodis, E. et al. A landscape of driver mutations in melanoma. Cell 150, 251–263 (2012).

87. Kichina, J. V, Rauth, S., Das Gupta, T. K. & Gudkov, A. V. Melanoma cells can tolerate high levels of transcriptionally active endogenous p53 but are sensitive to retrovirus-transduced p53. Oncogene 22, 4911–4917 (2003).

